# Machine Learning Predicts New Anti-CRISPR Proteins

**DOI:** 10.1101/854950

**Authors:** Simon Eitzinger, Amina Asif, Kyle E. Watters, Anthony T. Iavarone, Gavin J. Knott, Jennifer A. Doudna, Fayyaz ul Amir Afsar Minhas

**Author notes:** These authors contributed equally to this work (Simon Eitzinger, Amina Asif, Kyle E. Watters). To whom correspondence should be addressed., to 85 Extension: 3164. Department of Computer Science, University of Warwick, Coventry, CV4 7AL, UK.

## Abstract

The increasing use of CRISPR-Cas9 in medicine, agriculture and synthetic biology has accelerated the drive to discover new CRISPR-Cas inhibitors as potential mechanisms of control for gene editing applications. Many such anti-CRISPRs have been found in mobile genetic elements that disable the CRISPR-Cas adaptive immune system. However, comparing all currently known anti-CRISPRs does not reveal a shared set of properties that can be used for facile bioinformatic identification of new anti-CRISPR families. Here, we describe AcRanker, a machine learning based method for identifying new potential anti-CRISPRs directly from proteomes using protein sequence information only. Using a training set of known anti-CRISPRs, we built a model based on XGBoost ranking and extensively benchmarked it through non-redundant cross-validation and external validation. We then applied AcRanker to predict candidate anti-CRISPRs from self-targeting bacterial genomes and discovered two previously unknown anti-CRISPRs: AcrllA16 (ML1) and AcrIIA17 (ML8). We show that AcrIIA16 strongly inhibits *Streptococcus iniae* Cas9 (SinCas9) and weakly inhibits *Streptococcus pyogenes* Cas9 (SpyCas9). We also show that AcrIIA17 inhibits both SpyCas9 and SauCas9 with low potency. The addition of AcRanker to the anti-CRISPR discovery toolkit allows researchers to directly rank potential anti-CRISPR candidate genes for increased speed in testing and validation of new anti-CRISPRs. A web server implementation for AcRanker is available online at http://acranker.pythonanywhere.com/.

## INTRODUCTION

CRISPR-Cas systems use a combination of genetic memory and highly specific nucleases to form a powerful adaptive defense mechanism in bacteria and archaea (1–4). Due to their high degree of sequence specificity, CRISPR-Cas systems have been adapted for use as programmable DNA or RNA editing tools with novel applications in biotechnology, diagnostics, medicine, agriculture, and more (5–9). In 2013, the first anti-CRISPR proteins (Acrs) were discovered in *Pseudomonas aeruginosa* phages able to inhibit the CRISPR-Cas system (10). Since then, Acrs able to inhibit a wide variety of different CRISPR subtypes have been found (10–19). Multiple methods for identifying Acrs include screening for phages that escape CRISPR targeting (10, 19–23), guilt-by-association studies (12, 17, 24, 25), identification and screening of genomes containing self-targeting CRISPR arrays (11–13, 24), and metagenome DNA screening for inhibition activity (26, 27). Of these approaches, the ‘guilt-by-association’ search strategy is one of the most effective and direct, but it requires a known Acr to serve as a seed for the search. Thus, the discovery of one new validated Acr can lead to bioinformatic identification of others, as many Acrs have been discovered to be encoded in close physical proximity to each other, typically co-occurring in the same transcript with other Acrs or anti-CRISPR associated (aca) genes (12, 17). Screening approaches are particularly useful in this regard, as they can potentially identify new Acr families.

Identification of self-targeting CRISPR arrays can also help in predicting new Acr families. Typically, a CRISPR array with a spacer targeting the host genome (self-targeting) is lethal to the cell (28). However, if a mobile genetic element (MGE) present in the cell carries *acr* genes, the CRISPR-Cas system could be inhibited, and this may allow a cell with a self-targeting array to survive. To find new Acrs, genomes containing self-targeting arrays are identified through bioinformatic methods, and the MGEs within are screened for anti-CRISPR activity, eventually narrowing down to individual proteins (11, 24). Screens based on self-targeting also benefit from the knowledge of the exact CRISPR system that an inhibitor potentially exists for, as opposed to broad (meta-)genomic screens where a specific Cas protein has to be selected to screen against.

However, a weakness of all of these methods is that they are unable to predict *a priori* whether a gene may be an Acr, largely because Acr proteins do not share high sequence similarity or mechanisms of action (14, 16, 29–35). One theory to explain the high diversity of Acrs is the rapid mutation rate of the mobile genetic elements they are found in and the need to evolve with the co-evolving CRISPR-Cas systems trying to evade anti-CRISPR activity. Due to the relatively small size of most Acrs and their broad sequence diversity, simple sequence comparison methods for searching anti-CRISPR proteins are not expected to be effective. In this work, we report the development of AcRanker, a machine learning based method for direct identification of anti-CRISPR proteins. Using only amino acid composition features, AcRanker ranks a set of candidate proteins on their likelihood of being an anti-CRISPR protein. A rigorous cross-validation of the proposed scheme shows known Acrs are highly ranked out of proteomes. We then use AcRanker to predict 10 new candidate Acrs from proteomes of bacteria with self-targeting CRISPR arrays and biochemically validated three of them. Our machine learning approach presents a new tool to directly identify potential Acrs for biochemical validation using protein sequence alone.

## MATERIALS AND METHODS

### Data collection and preprocessing

To model the task of anti-CRISPR protein identification as a machine learning problem, a dataset consisting of examples from both positive (anti-CRISPR) and negative (non-anti-CRISPR) classes was needed. We collected anti-CRISPR information for proteins from the Anti-CRISPRdb (36). The database contained information for 432 anti-CRISPR proteins. We used CD-HIT to identify a non-redundant set (at 40% sequence similarity threshold) of 20 experimentally verified Acrs (Table S1) (37). These proteins belong to different Acr classes: 12 of the proteins are active against class I-F CRISPR Cas systems, 4 against I-E and 4 against II-A (10, 13, 17, 20, 22). This set constitutes positive class of our dataset. We downloaded complete proteomes of source species to which each of these proteins belong. Proteins in these proteomes with <40% sequence similarity with the set of known anti-CRISPR proteins were used to construct the negative dataset. For independent testing of the method, a dataset comprising recently found Acrs (12) was used (Table S2). The proteins used for development of the machine learning model in AcRanker belonged to classes I-F, I-E, and II-A (36), so only the Acrs that belonged to either of these classes were chosen. Source proteomes for all these proteins were downloaded.

### Feature Extraction

In line with existing machine learning based protein function prediction techniques, we used sequence features (38) based on amino acid composition and grouped dimer and trimer frequency counts (39). For this purpose, amino acids are first grouped into seven classes based on their physicochemical properties (39) (Table S3) and the frequency counts of all possible groups labeled as dimers and trimers in a given protein sequence are used in conjunction with amino acid composition. All three types of features (amino acid composition, di- and tri- meric frequency counts) are normalized to unit norm resulting in a 20 + 7^2^ + 7^3^ = 412-dimensional feature vector representation for a given protein sequence (40, 41).

### Machine learning model

The underlying machine learning model for AcRanker has been built using EXtreme Gradient Boosting (XGBoost) (42). In machine learning, boosting is a technique in which multiple weak classifiers are combined to produce a strong classifier (42). XGBoost is a tree based method (42) that uses boosting in an end-to-end fashion, i.e., every next tree tries to minimize the error produced by its predecessor. XGBoost has been shown to be a fast and scalable learning algorithm and has been widely used in many machine learning applications.

In this work, we have used XGBoost as a pairwise ranking model to rank constituent proteins in a given proteome in descending order of their expected Acr behavior. The XGBoost model is trained in a species-specific manner to produce higher scores for anti-CRISPR proteins as compared to non-anti-CRISPR proteins in a given proteome. In comparison to conventional XGBoost classification, the pairwise ranking model performed better in terms of correctly identifying known anti-CRISPR proteins in test proteomes in cross-validation (comparison not shown for brevity). Specifically, given a set of training proteomes *S* each with one or more known anti-CRISPR proteins, our objective is to obtain an XGBoost predictor *f*(***x***; *θ*) with learnable parameters *θ* that generates a prediction score for a given protein sequence represented in terms of its feature vector ***x***. In training, we require the model to learn optimal parameters *θ*^*^ such that the score *f*(***p***; *θ*^*^) for a positive example ***p*** (known Anti-CRISPR protein) should be higher than *f*(***n***; *θ*^*^) for all negative examples ***n*** (non-Anti-CRISPR proteins) within the same species. The hyperparameters of the learning model are selected through cross validation and optimal results are obtained with: number of estimators set at 120, learning rate of 0.1, subsampling of 0.6 and maximum tree depth of 3.

### Performance Evaluation

To evaluate the performance of the machine learning model, we have performed leave-one-proteome-out cross-validation as well as validation over an independent test set. In a single fold of leave-one-proteome-out cross-validation, we set aside the source proteome of a given anti-CRISPR protein for testing and train on all other proteomes. To ensure an unbiased evaluation, all sequences in the training set with a sequence identity of 40% or higher with any test protein or among themselves are removed from the training set. Furthermore, all proteins in the test set with more than 40% sequence identity with known anti-CRISPR proteins in the training set are also removed. This ensures that there is only one known anti-CRISPR protein in the test set in a single fold. The XGBoost ranking model is then trained and the prediction scores for all proteins in the test set are computed. Ideally, the known anti-CRISPR protein in the proteome should score the highest across all proteins in the given test proteome. This process is then repeated for all proteomes in our dataset. The rank of the known anti-CRISPR protein in its source proteome is used as a performance metric.

In bacteria, Acrs are usually located within prophage regions (13, 43). Based on this premise, in another experiment for model evaluation, we passed only the proteins found within prophage regions to the model. To identify the prophage regions for a given bacterial proteome we used PHASTER (PHAge Search Tool Enhanced Release) web server (44) which accepts a bacterial genome and annotates prophage regions in it. The decision scores are computed for all phage proteins identified by PHASTER in the test proteome.

As a baseline for comparison in leave-one-out cross-validation, BLAST (Basic Local Alignment Search Tool) (45) similarity was used. For each protein in a given test proteome, we compute BLASTp scores with the set of known Acrs and rank proteins in the increasing order of the respective e-values.

For independent validation, the ranking based XGBoost model trained over sequence features for all 20 source proteomes (Table S1) has been tested for recently discovered Acrs (Table S2) by Marino et al. (12) which are not part of our training set. The rank of known Acr in its corresponding proteome was computed. Here again, we evaluated the model for both complete proteomes and respective MGE subset identified by PHASTER.

### AcRanker Webserver

A webserver implementation of AcRanker is publicly available at http://acranker.pythonanywhere.com/. The webserver accepts a proteome file in FASTA format and returns a ranked list of proteins. The Python code for the webserver implementation is available at the URL: https://github.com/amina01/AcRanker.

### Acr candidate selection

Self-targeting Spacer Searcher (STSS; https://github.com/kew222/Self-Targeting-Spacer-Searcher) (11) was run with default parameters using ‘Streptococcus’ as a search term for the NCBI genomes database, which returned a list of all self-targets found in those genomes. Whether known Acr genes were present in each of the self-targeting genomes was checked using a simple blastp search using default parameters with the Acr proteins stored within STSS. Twenty self-targeting genomes that contained at least one self-target with a 3′-NRG PAM were chosen for further analysis with AcRanker. Prophage regions were predicted using PHASTER (44). Proteins within the prophage regions were ranked with AcRanker.

To select individual gene candidates for synthesis and biochemical validation, the six highest ranked proteins from each genome were examined by visual inspection for a strong promoter, a strong ribosome binding site, and an intrinsic terminator. Promoters were searched for manually by looking for sequences closely matching the strong consensus promoter sequence TTGACA-17(+/−1)N-TATAAT upstream of the Acr candidate gene, or any genes immediately preceding it. The presence of a strong ribosome binding site (resembling AGGAGG) near the start codon was similarly searched for and was required to be upstream of a gene candidate for selection. Last, given the nature of Acrs to be clustered together, genes neighboring the best candidates were also selected for further testing/validation.

### Protein expression and purification

Each of the Acr candidates (Table S4) were cloned into a custom vector (pET-based expression vector) such that each protein was N-terminally tagged with a 10xHis sequence, superfolder GFP, and a tobacco etch virus (TEV) protease cleavage site, available on Addgene. Each Cas effector (Table S5), *Acidaminococcus sp*. Cas12a (AsCas12a), *Streptococcus pyogenes* Cas9 (SpyCas9), *Staphylococcus aureus* Cas9 (SauCas9) and *Streptococcus iniae* Cas9 (SinCas9), were expressed as N-terminal MBP fusions. Proteins were produced and purified as previously described (32). Briefly, *E. coli* Rosetta2 (DE3) containing Acr or Cas9 expression plasmids were grown in Terrific Broth (100 μg/mL ampicillin) to an OD_600_ of 0.6-0.8, cooled on ice, induced with 0.5 mM isopropyl-b-D-thiogalactoside and incubated with shaking at 16°C for 16 h. Cells were harvested by centrifugation, resuspended in wash buffer (20 mM Tris-Cl (pH 7.5), 500 mM NaCl, 1 mM tris(2-carboxyethyl)phosphine (TCEP), 5% (v/v) glycerol) supplemented with 0.5 mM phenylmethanesulfonyl fluoride and cOmplete protease inhibitor (Roche), lysed by sonication, clarified by centrifugation and purified over Ni-NTA Superflow resin (Qiagen) in wash buffer supplemented with 10 mM (wash) or 300 mM imidazole (elution). Elution fractions were pooled and digested overnight with recombinantly expressed TEV protease while dialysed against dialysis buffer (20 mM Tris-Cl (pH 7.5), 125 mM NaCl, 1 mM TCEP, 5% (v/v) glycerol) at 4°C. The cleaved proteins were loaded onto an MBP-Trap (GE Healthcare) upstream of a Heparin Hi-Trap (GE Healthcare) in the case of SpyCas9, SauCas9 and SinCas9. Depending on the pI, TEV digested Acrs were loaded onto a Q (ML1, ML2, ML3, ML6, ML8 and ML10), heparin (ML4, ML5), or SP (ML7 and ML9) Hi-Trap column. Proteins were eluted over a salt gradient (20 mM Tris-Cl (pH 7.5), 1 mM TCEP, 5% (v/v) glycerol, 125 mM – 1 M KCl). The eluted proteins were concentrated and loaded onto a Superdex S200 Increase 10/300 (GE Healthcare) for SpyCas9, SauCas9, SinCas9 or Superdex S75 Increase 10/300 (GE Healthcare) for all the Acr candidates developed in gel filtration buffer (20 mM HEPES-K (pH 7.5), 200 mM KCl, 1 mM TCEP and 5% (v/v) glycerol). The absorbance at 280 nm was measured by Nanodrop and the concentration was determined using an extinction coefficient estimated based on the primary amino acid sequence of each protein. Purified proteins were concentrated to approximately 50 μM for Cas9 effectors and 100 μM for Acr candidates. Proteins were then snap-frozen in liquid nitrogen for storage at −80 °C. Purity and integrity of proteins was assessed by 4-20% gradient SDS-PAGE (Coomassie blue staining, Figure S1A) and LC-MS (Figure S1B).

### RNA preparation

All RNAs (Table S6) were transcribed *in vitro* using recombinant T7 RNA polymerase and purified by gel extraction as described previously (46). Briefly, 100 μg/mL T7 polymerase, 1 μg/mL pyrophosphatase (Roche), 800 units RNase inhibitor, 5 mM ATP, 5 mM CTP, 5 mM GTP, 5 mM UTP, 10 mM DTT, were incubated with DNA target in transcription buffer (30 mM Tris-Cl pH 8.1, 25 mM MgCl_2_, 0.01% Triton X-100, 2 mM spermidine) and incubated overnight at 37**°**C. The reaction was quenched by adding 5 units RNase-free DNase (Promega). Transcription reactions were purified by 12.5% (v/v) urea-denaturing PAGE (0.5x Tris-borate-EDTA (TBE)) and ethanol precipitation.

### *In vitro* cleavage assay

*In vitro* cleavage assays were performed at 37°C in 1X cleavage buffer (20 mM Tris-HCl pH 7.5, 100 mM KCl, 5 mM MgCl_2_, 1 mM DTT and 5% glycerol (v/v)) targeting a PCR amplified fragment of double-stranded DNA (Table S7). For all cleavage reactions, the sgRNA was first incubated at 95°C for 5 min and cooled down to room temperature. The Cas effectors (SpyCas9, SauCas9, AsCas12a at 100 nM and SinCas9 at 200 nM respectively) were incubated with each candidate Acr protein at 37°C for 10 min before the addition of sgRNA (SpyCas9, SauCas9, AsCas12a sgRNA at 160 nM and SinCas9 sgRNA at 320 nM respectively) to form the RNP at 37°C for 10 min. The DNA cleavage reaction was then initiated with the addition of DNA target and reactions incubated for 30 min at 37°C before quenching in 1X quench buffer (5% glycerol, 0.2% SDS, 50 mM EDTA). Samples were then directly loaded to a 1% (w/v) agarose gel stained with SYBRGold (ThermoFisher) and imaged with a BioRad ChemiDoc.

### Competition binding experiment

The reconstitution of the SinCas9-sgRNA-ML1 and SinCas9-sgRNA-AcrIIA2 complex was carried out as previously described (47). Briefly, purified SinCas9 and *in vitro* transcribed sgRNA were incubated in a 1:1.6 molar ratio at 37°C for 10 min to form the RNP. To form the inhibitor bound complexes, a 10-fold molar excess of AcrIIA16 or AcrIIA2 were added and incubated with the RNP complex at 37°C for 10 min. For the competition binding experiment, a 10-fold molar excess of AcrIIA16 was first incubated with the RNP complex at 37°C before incubation with a 10-fold molar excess of AcrIIA2 at 37°C for 10 min. Each complex was then purified by analytical size-exclusion chromatography (Superdex S200 Increase 10/300 GL column, GE Healthcare) pre-equilibrated with the gel filtration buffer (20 mM HEPES-K (pH 7.5), 200 mM KCl, 1 mM TCEP and 5% (v/v) glycerol) containing 1 mM MgCl_2_. The peak fractions were concentrated by spin concentration (3-kDa cutoff, Merck Millipore), quenched in 1X SDS-Loading dye (2% w/v SDS, 0.1% w/v bromophenol blue and 10% v/v glycerol) and boiled down to 20 μl before loading onto a 4-20% gradient SDS-PAGE.

### Mass spectrometry

Protein samples were analyzed using a Synapt mass spectrometer as described elsewhere (48).

## RESULTS

### Cross-validation by single proteome omission

In this work, we have developed AcRanker, a machine learning model that accepts a proteome as input and ranks its constituent proteins in decreasing order of their expected Acr character. We have used EXtreme Gradient Boosting (XGBoost) based ranking (42) with 1, 2 and 3-mer amino acid composition as input features (38) to train on a dataset comprised of 20 experimentally verified Acrs taken from the anti-CRISPRdb (29, 36) (Table S1) and their source proteomes. To evaluate the performance of AcRanker, we performed leave-one-out cross-validation as well as testing over an independent set of proteins. Out of the 20 known Acr proteomes tested individually, we observed that the ranking-based model ranked seven Acrs higher than other proteins in their respective proteomes (Table 1). In total, 14 out of the 20 known Acrs are ranked within the top 5% in their respective proteomes (Table 1).

**Table 1.** Results for leave-one-out cross-validation. Each row of the table indicates which Acr was excluded from the training dataset and used as a test dataset, and each number displayed is the ranking of the known Acr received from the indicated test proteome using either the blastp search against all other known Acrs (BLAST) or AcRanker. The Acrs from bacterial proteomes - AcrIF6, AcrIF9, AcrIF10, AcrIIA1, AcrIIA2, and AcrIIA4 - were also ranked using only the subset of proteins predicted to reside within prophages as predicted by PHASTER (44). Two Acrs from bacterial proteomes did not occur in the predicted prophages and are indicated by dash placeholders. Prophage proteome subset fields have been left empty for Acrs from phage proteomes.

| Accession No. | Anti-CRISPR family | Complete Proteome |  |  | Prophage Subset |  |
| --- | --- | --- | --- | --- | --- | --- |
|  |  | Proteome Size | BLAST rank | AcRanker rank | Proteome Size | AcRanker rank |
| YP_007392738.1 | AcrIE1 | 57 | 33 | **1** | - | - |
| YP_007392439.1 | AcrIE2 | 54 | 18 | 2 | - | - |
| YP_950454.1 | AcrIE3 | 52 | 17 | **1** | - | - |
| NP_938238.1 | AcrIE4 | 54 | 11 | **1** | - | - |
| YP_007392342.1 | AcrIF1 | 56 | 21 | 11 | - | - |
| YP_002332454.1 | AcrIF2 | 51 | 34 | **1** | - | - |
| YP_007392440.1 | AcrIF3 | 54 | 5 | **1** | - | - |
| YP_007392799.1 | AcrIF4 | 57 | 36 | 3 | - | - |
| YP_007392740.1 | AcrIF5 | 57 | 26 | 19 | - | - |
| WP_043884810.1 | AcrIF6 | 6095 | **1** | 80 | 361 | 15 |
| WP_019933870.1 | AcrIF6 | 3045 | **1** | 13 | 72 | **1** |
| WP_014702809.1 | AcrIF6 | 2689 | **1** | 130 | 57 | - |
| ACD38920.1 | AcrIF7 | 57 | 20 | **1** | - | - |
| AFC22483.1 | AcrIF8 | 68 | 30 | **1** | - | - |
| WP_031500045.1 | AcrIF9 | 4928 | 198 | 333 | 37 | - |
| KEK29119.1 | AcrIF10 | 3552 | 189 | 17 | 70 | 2 |
| AEO04364.1 | AcrIIA1 | 2951 | 183 | 770 | 146 | 87 |
| AEO04363.1 | AcrIIA2 | 2952 | 210 | 16 | 146 | 3 |
| AEO04689.1 | AcrIIA4 | 2951 | 59 | 21 | 146 | 4 |
| D4276_028 | AcrIIA5 | 54 | 5 | 8 | - | - |

Generally, we observe that the machine learning rankings for Acrs contained in phage proteomes are much better than those contained in bacterial proteomes, likely due to their smaller size (Table 1). To test if the relative rankings of the known Acrs found within bacterial proteomes would improve in the context of only prophage-derived proteins, we identified which proteins in the bacterial proteomes were found within prophages using PHASTER (44) and used only that subset to test both models. With the prophage subsets we did observe a higher ranking for the known Acrs due to the removal of higher-ranking proteins not found in the predicted prophages (Table 1).

As a baseline, we also compared the rankings obtained from the machine learning model to a blastp (45) comparison (Table 1). For each excluded Acr in the leave-one-out train/test cycles, the excluded Acr’s proteome was used as a query set to BLAST against the 19 other Acrs used for training and the resulting e-values ranked from lowest to highest. The BLAST search method, however, only returned the highest rank for the AcrIF6 family, likely because three distant homologs (using the <40% identity threshold) were included in the training dataset. Interestingly, we also observed that the BLAST method gave a higher rank than AcRanker for AcrIIA1, which contains a motif (helix-turn-helix) that is found in some other known Acrs (13, 21, 24, 25). The rankings of all other Acrs fell outside of the top 5%, demonstrating the diversity of Acr families and the difficulty of predicting new Acrs *de novo*.

### Independent set validation

To validate AcRanker, we used an independent testing dataset of 10 recently discovered Acrs not a part of the training dataset (Table S2) (49). Of these 10 Acrs, one is found in a phage (AcrIF14) and four (AcrIE4-F7, AcrIF11, AcrIF11.1, and AcrIF11.2) were predicted to be in a prophage region using PHASTER. For the proteins predicted to be in a prophage both the complete bacterial and phage proteome was ranked with AcRanker, otherwise only the complete proteome was ranked (Table S8). The results from the complete bacterial proteomes did not perform well (Table S8), with AcrIE5 and AcrIF12 receiving ranks within the top 10. However, of the five proteins found within a phage/prophage, AcRanker ranked three within the top five, including one with the highest rank (Table 2).

**Table 2.**
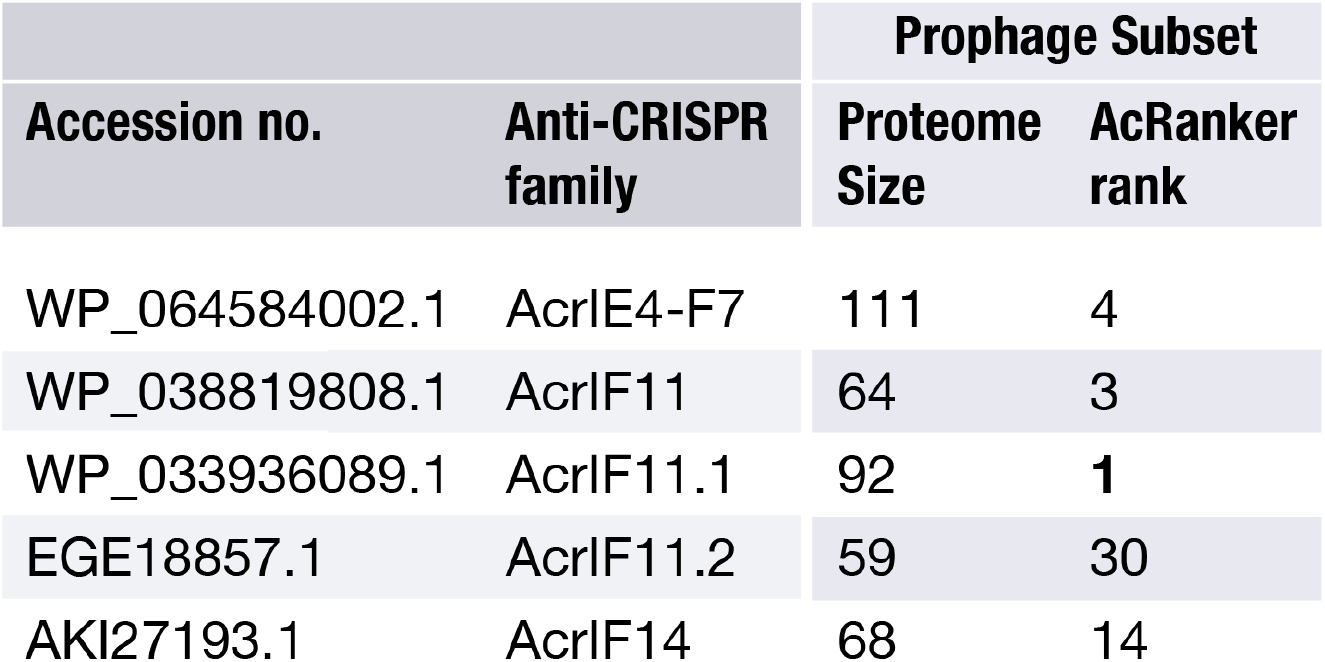
Independent testing set validation results. Five proteomes containing non-redundant (<40% sequence identity) Acrs from bacterial proteomes that had Acrs within PHASTER-predicted prophages were tested with AcRanker.

### anti-CRISPR candidate selection

Encouraged by the number of highly ranked Acrs from the test dataset, we proceeded to apply AcRanker to predict novel anti-CRISPRs from self-targeting genomes. Given the ubiquity of *Streptococcus pyogenes* Cas9 (SpyCas9) in gene editing and our inclusion of known SpyCas9 Acrs in the machine learning training dataset (AcrIIA1, AcrIIA2, AcrIIA4, AcrIIA5), we chose to focus specifically on *Streptococcus* species containing Cas9 proteins homologous to SpyCas9.

We began by generating a list of *Streptococcus* genomes containing at least one self-targeting type II-A CRISPR system using Self-Target Spacer Searcher, which has been previously described (11). We found 385 instances of self-targeting from type II-A CRISPR arrays occurring within 241 *Streptococcus* genome assemblies, six of which contained known Acrs. Of these 241 self-targeting arrays, we looked for instances where the target sequence was flanked by the 3′ NRG protospacer adjacent motif (PAM) characteristic of SpyCas9 and observed that it was present in 20 genomes. These 20 self-targeting arrays would be expected to be lethal for close homologs of SpyCas9, suggesting that other factors, such as the presence of Acrs (11), are preventing CRISPR self-targeting and cell death (Table S9). During our original search of these 20 genomes, *Streptococcus iniae* strain UEL-Si1 was the only one that contained a previously discovered Acr, AcrIIA3 (13), providing a large proteome space to search for novel *acr* genes.

To identify new *acr* gene candidates, we first used PHASTER (44) to predict all of the prophages residing within the 20 self-targeting *Streptococcus* genomes as well as an additional *Listeria monocytogenes* genome (strain R2-502) containing a type II-A self-targeting CRISPR system (with six self-targets) and three well-known AcrIIA genes (13) We included the *Listeria* strain to determine if the known Acrs within it were returned as the top ranked genes, and if not, test the higher ranking genes as potential additional Acrs within a known Acr-harboring strain. We created lists of the annotated proteins found within each genome’s set of prophages. These proteins lists were then ranked with AcRanker to predict the 10 highest ranked genes most likely to be an *acr* (Table S10). Of the approximately 200 genes returned, a subset was selected as the most likely to be undiscovered *acr* genes for further biochemical testing, based on previous observations that many Acrs are: 1) encoded in operons along other *acrs* 2) typically short genes, and 3) often have transcripts driven by strong promoters and ribosome binding sites that frequently end with intrinsic terminator sequences (11, 13, 24) (Figure 1).

**Figure 1.**
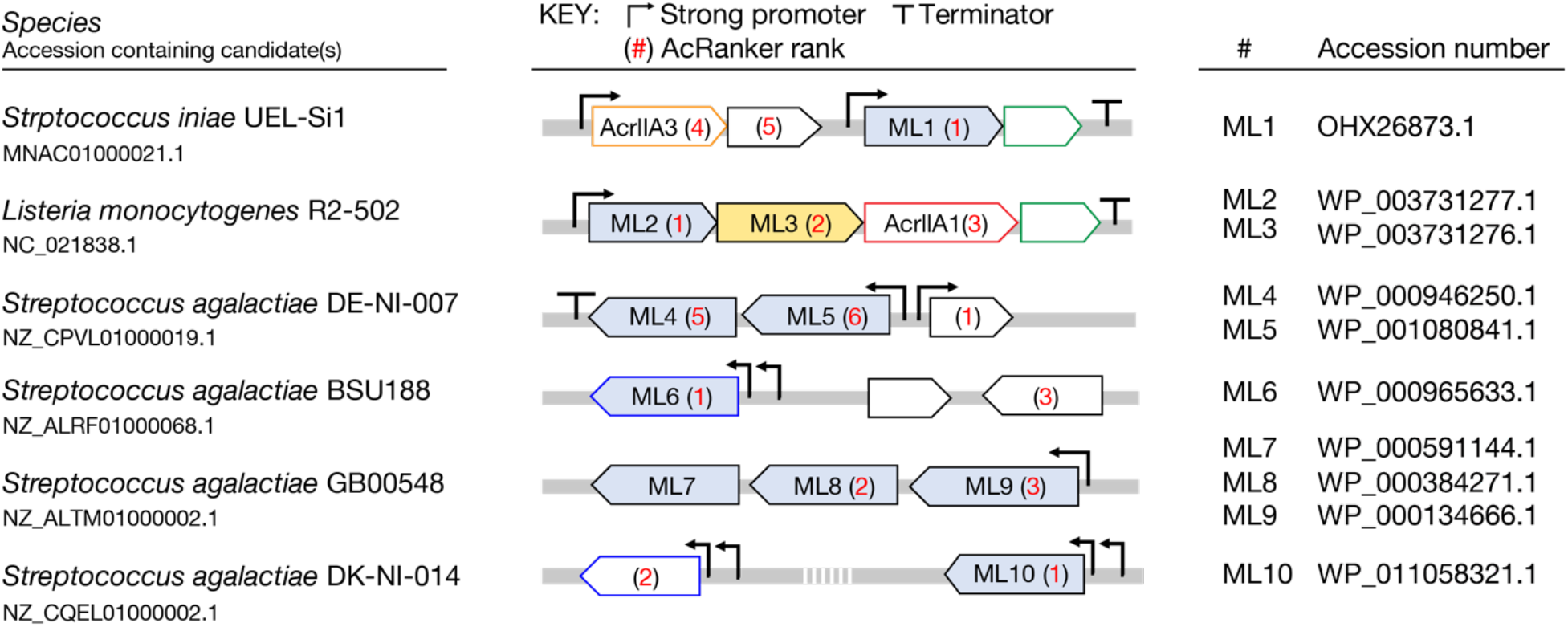
Acr candidates selected for biochemical testing. Ten Acr candidates were selected from manual inspection for further biochemical testing (blue). Each candidate is shown in its genomic context with its assigned rank from AcRanker noted in red. Homologous proteins share the same color border (green, blue). Homologs of AcrIIA3 (orange border) and AcrIIA1 (red border) are indicated. While testing the ML candidates, ML3 (yellow) has been identified as a specific inhibitor of LmoCas9 (25).

As with the previous testing dataset, we observed that the known *acr* genes were highly ranked within the test proteomes. Interestingly, other proteins contained in the same, or overlapping, transcripts as the known Acrs ranked higher with AcRanker (ML1 and ML2). We took these candidates as well as eight others (ML3-ML10) containing the features described above (Figure 1).

### Biochemical validation of novel Acrs identified by AcRanker

To determine if the identified proteins were inhibitors of SpyCas9, we purified each candidate and tested their ability to directly inhibit DNA targeting *in vitro*. Of the ten candidate inhibitors, nine were successfully cloned, expressed and purified (Figure S1A and B). To assess inhibition of DNA targeting *in vitro*, we first assayed the ability of SpyCas9 to cleave double stranded (ds) DNA when incubated in the presence of a 50-fold excess of each candidate Acr (Figure 2A). While SpyCas9 was capable of complete DNA target cleavage, the generation of DNA cleavage products was attenuated in the presence of the positive control inhibitor AcrIIA4 and the candidates ML1 or ML8. To determine the potency of inhibition, we tested the ability of SpyCas9 to cleave the DNA target in the presence of a dilution series of ML1 or ML8 (Figure 2B). In contrast to AcrIIA4, an established potent inhibitor of SpyCas9 (13), both ML1 and ML8 inhibited SpyCas9 with around a 10-fold lower potency. We wondered if the high concentration of ML1 or ML8 required to completely inhibit Cas9 might represent an *in vitro* concentration-dependent artefact. To explore this, we assayed SpyCas9 DNA cleavage against a titration series of either non-target DNA competitor, BSA, ML2, or ML3 and observed no significant inhibition of SpyCas9, even with a 100-fold excess (Figure S2B-D). Taken together, these data indicated that both ML1 and ML8 weakly inhibit SpyCas9 DNA cleavage *in vitro*.

**Figure 2.**
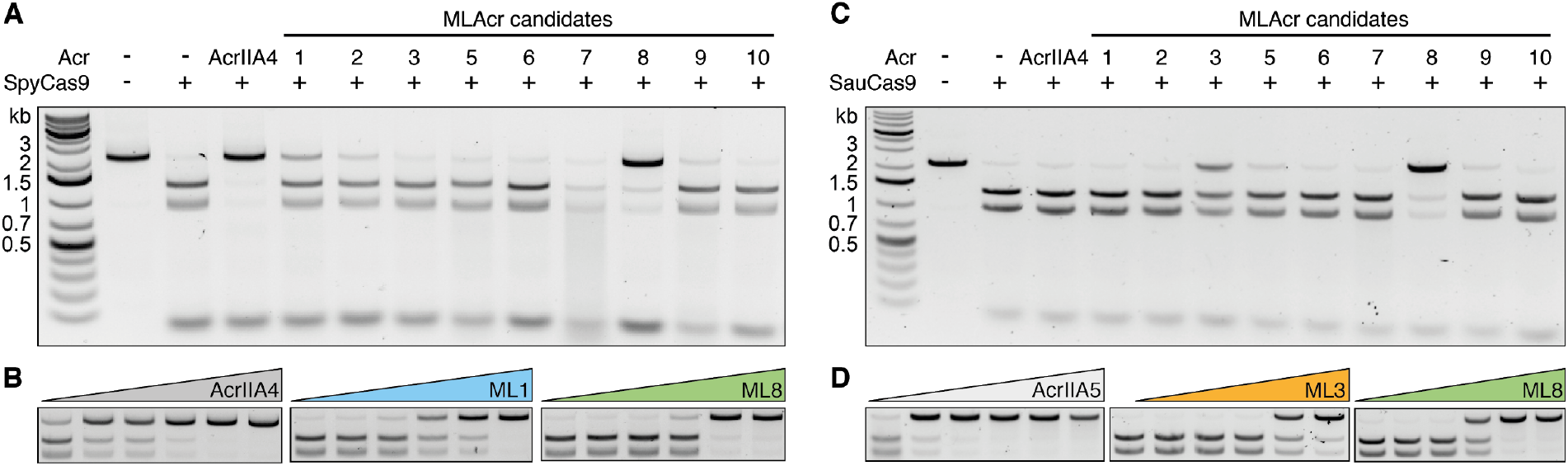
Inhibition of SpyCas9 and SauCas9 by newly discovered Acr candidates. **(A)** *In vitro* cleavage of dsDNA by SpyCas9 in the absence or presence of a 50-fold excess of AcrIIA4 (positive control) and each Acr candidate. (**B**) *In vitro* cleavage of dsDNA by SpyCas9 in the presence of increasing concentrations of (left to right) AcrIIA4 (positive control), ML1 and ML8 (Acr:RNP 0.1-, 1-, 2-,10-, 50- and 100-fold excess from left to right). (**C**) *In vitro* cleavage of dsDNA by SauCas9 in the absence or presence of a 25-fold excess of each Acr candidate. (**D**) *In vitro* cleavage of dsDNA by SauCas9 in the presence of increasing concentrations of (left to right) AcrllA5 (positive control, Acr:RNP 0.1-, 1-, 2-,4-, 8- and 10-fold excess from left to right), ML3 and ML8 (Acr:RNP 0.1-, 1-, 2-,10-, 50- and 100-fold excess from left to right). Uncropped gel images for panels B and D are shown in Figure S2 and S3.

We next tested the ability of the AcRanker-generated candidates to inhibit *Staphylococcus aureus* (SauCas9), another Cas9 commonly used for gene editing (50, 51) to determine whether any of the candidates identified from self-targeting *Streptococcus* genomes had broader Cas9 inhibition activity. At a 25-fold excess relative to the SauCas9 RNP complex, ML3 and ML8 were able to inhibit SauCas9 dsDNA cleavage (Figure 2C). To determine potency, we incubated a dilution series of either ML3 or ML8 with SauCas9 before the addition of the DNA target. However, in comparison to AcrIIA5, an established strong inhibitor of SauCas9 (20, 24), both Acr candidates inhibited SauCas9 with approximately 50-fold lower potency (Figure 2D, Figure S3A and B), an activity we confirmed was not due to a false positive from the high concentration of protein in the assay (Figure 3A).

**Figure 3.**
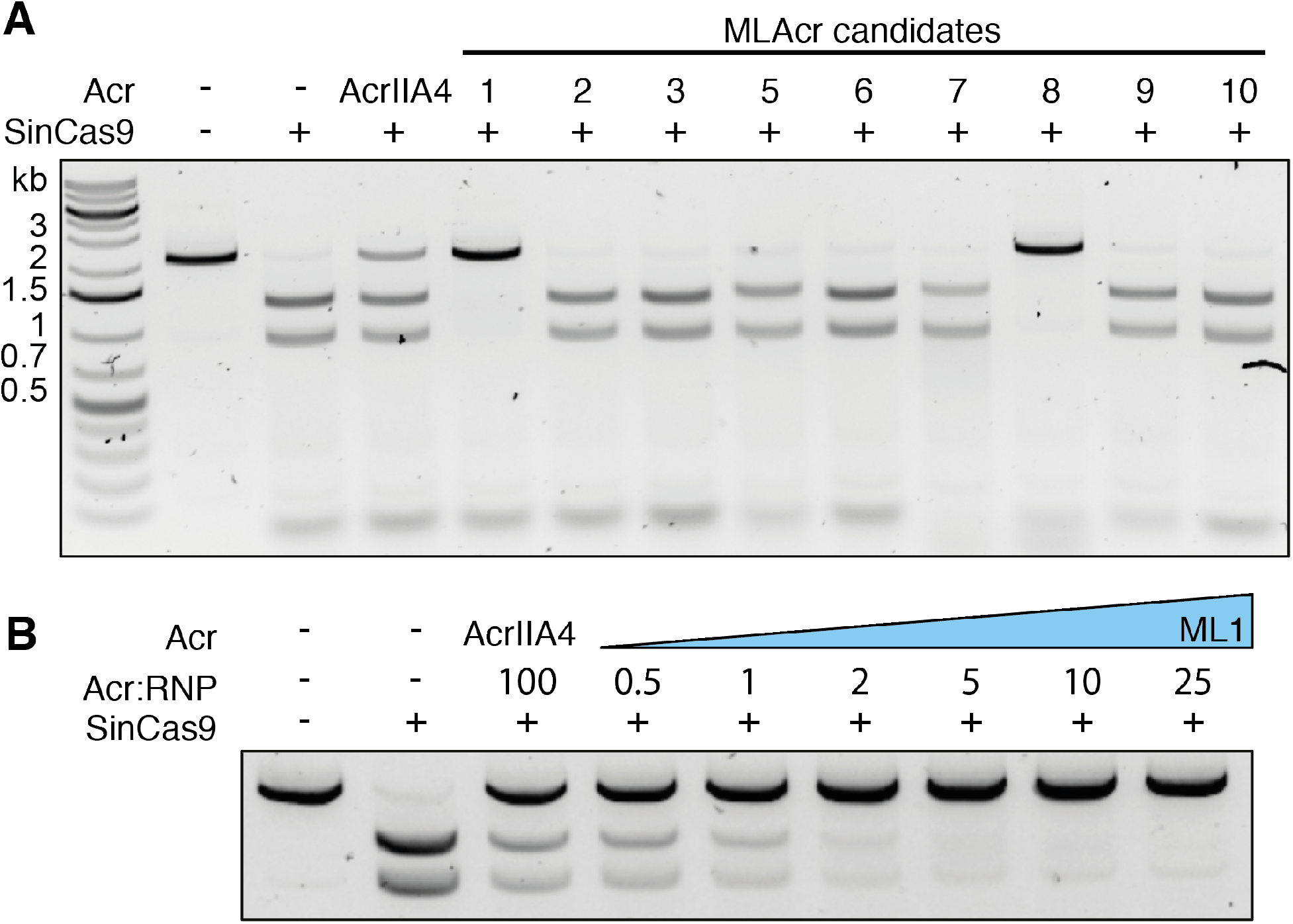
ML1 and ML8 inhibit SinCas9 with ML1 showing very high potency. **(A)** *In vitro* cleavage of dsDNA by SinCas9 in the absence or presence of a 50-fold excess of each Acr candidate. (**B**) *In vitro* cleavage of dsDNA by SinCas9 in the presence of increasing concentrations of ML1. The uncropped gel image for panel B is shown in Figure S5.

Given the relatively weak inhibition of both SpyCas9 and SauCas9, we next tested the specificity of ML1, ML3 and ML8 by assaying their ability to block DNA targeting by either AsCas12a or the restriction enzyme AlwNI. Neither AcrIIA4, ML1, ML3, nor ML8 were able to inhibit DNA targeting by AlwNI, consistent with them being specific inhibitors of CRISPR effectors (Figure S4A and B). Consistent with this, inhibition of AsCas12a was only observed with ML1 and ML8 at a 100-fold excess (Figure S4C). Taken together, our data are consistent with ML1, ML3, and ML8 being low potency inhibitors of SpyCas9 (ML1 and ML8) or SauCas9 (ML3 and ML8). Interestingly, while testing ML1-ML10 for Acr activity, Osuna, et al. described AcrIIA12, a specific inhibitor of LmoCas9 in plaque assays, which shares the same sequence as ML3 (25).

### ML1: a potent inhibitor of SinCas9

ML1 was identified in the *Streptococcus iniae* (Sin) genome. Previous studies have reported anti-CRISPRs can exhibit either selective or broad-spectrum inhibition of divergent Cas effectors (14, 32). Given that SinCas9 is 70.10% identical to SpyCas9 and only 25.58% identical to SauCas9 we wondered if ML1 might be a more potent inhibitor of SinCas9. To explore this, we cloned, expressed, and purified SinCas9 protein for use in *in vitro* DNA targeting assays. Like SpyCas9, SinCas9 was capable of cleaving dsDNA targets proximal to an NGG PAM using a sgRNA derived from a fusion of the tracrRNA and crRNA (Figure 3A, Figure S6). Similar to SpyCas9, both ML1 and ML8 inhibited DNA cleavage by SinCas9. Using a titration of ML1, we again assayed the potency of SinCas9 inhibition (Figure 3B, Figure S5B). Strikingly, in contrast to the weak inhibition of SpyCas9, ML1 was able to potently inhibit DNA cleavage by SinCas9 (Figure 3B). To investigate at which step ML1 inactivates SinCas9 function, we carried out *in vitro* cleavage assays where ML1 was incubated with SinCas9 before and after the addition of sgRNA (Figure S5C). In both cases the DNA cleavage activity of SinCas9 was potently inhibited, suggesting that ML1 inhibits activity after sgRNA binding to Cas9.

A number of reported type-IIA Acrs inhibit their cognate Cas9 by competing with target DNA through PAM mimicry (47, 52). We noted that SinCas9 was susceptible to inhibition by AcrIIA4 (Figure 3A) and AcrIIA2 (Figure S5D), both PAM mimics that inhibit PAM recognition by SpyCas9 (15, 47). Like these established PAM mimics, ML1 is a small protein with a predicted negatively charged surface potential (isoelectric point of 4.3), suggesting that it too might compete with target DNA. To explore this idea, we developed a competition binding experiment to assay if the association of ML1 with SinCas9 might prevent the binding of AcrIIA2 (Figure 4A). Firstly, we incubated either AcrIIA2 or ML1 with the SinCas9-sgRNA complex and observed a stable SinCas9-sgRNA-Acr complex on a gel filtration column (Figure 4B, Figure S7A) with the complex components all resolvable on a protein gel (Figure 4C, Figure S7B). To determine if ML1 binding to the SinCas9 RNP could prevent AcrIIA2 binding, we first formed the SinCas9-sgRNA-ML1 complex and then incubated with AcrIIA2 before resolving over a column. Incubating ML1 with the SinCas9 RNP before adding AcrIIA2 abolished AcrIIA2 co-elution with SinCas9-sgRNA (Figure 4C, Figure S7B), suggesting that ML1 might occupy the same site on SinCas9. Collectively, these data are consistent with a model where ML1 directly binds to the SinCas9-sgRNA complex to form a complex that is incompatible with AcrIIA2 binding.

**Figure 4.**
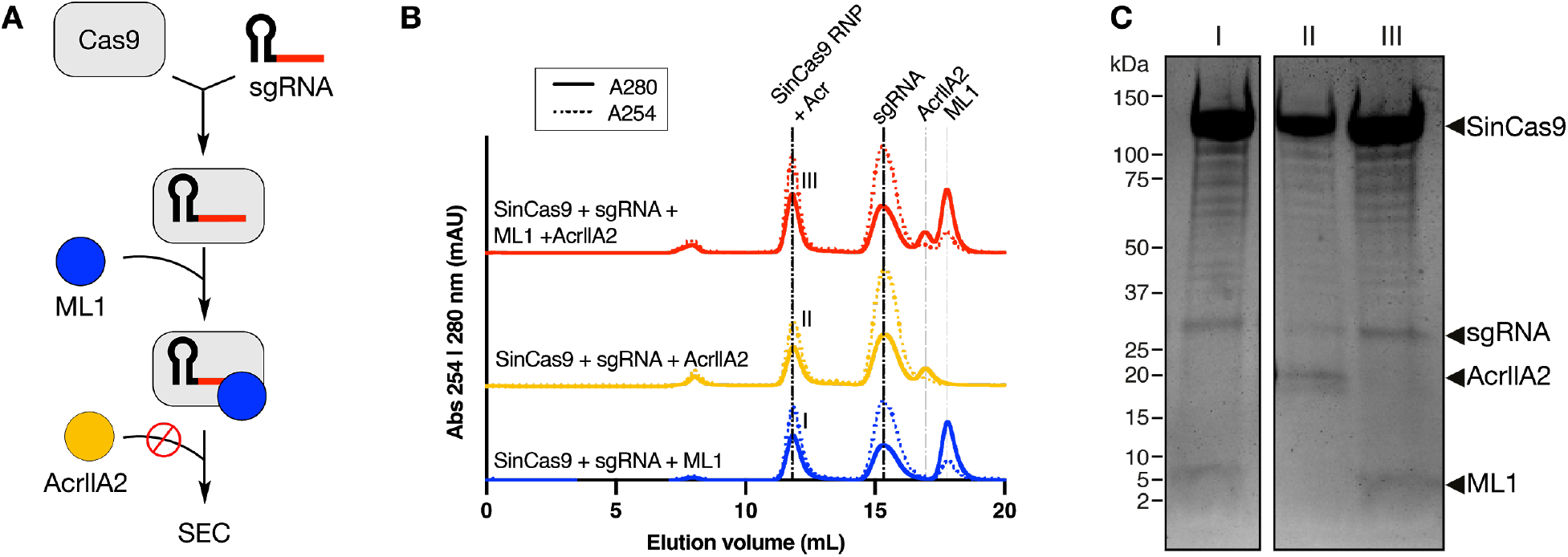
ML1 competes with AcrIIA2 to bind to the SinCas9-sgRNA complex. **(A)** Flowchart for the competition binding experiment between ML1 and AcrIIA2. Binding of the Acr to the SinCas9-sgRNA RNP was reconstituted using size-exclusion chromatography (SEC). (**B**) Size-exclusion chromatogram of SinCas9-sgRNA in the presence of either ML1, AcrIIA2 or both Acrs with AcrIIA2 added after ML1. (**C**) Coomassie-stained polyacrylamide gel illustrating the components of the SinCas9-RNP fraction annotated (I), (II), and (III) in panel B.

## DISCUSSION

With the growth of the anti-CRISPR field, there has been a need for improved tools to search the extensive proteomic space to find new anti-CRISPRs more efficiently. In this work we developed a machine learning method, AcRanker, which allows for direct prediction of Acr genes *de novo* with high accuracy and minimal knowledge *a priori.* Using a combination of AcRanker and self-targeting information from STSS (11), we were able to quickly reduce to a few top Acr gene candidates for direct synthesis and testing of anti-CRISPR properties. We identified two novel Acrs: here named AcrIIA16 and AcrIIA17. AcrIIA16 (ML1) inhibits *Streptococcus iniae* Cas9 (SinCas9) with high potency and *Streptococcus pyogenes* Cas9 (SpyCas9) with low potency. With only 64 amino acids and a molecular weight of 7.3 kDa, to our knowledge it is the smallest type II Acr found to date. Based on the negative charge of AcrIIA16 and its competitive binding with AcrIIA2, we speculate that AcrIIA16 inhibits Cas9 dsDNA cleavage via a similar mechanisms of PAM mimicry. In addition, we found AcrIIA17 (ML8), a broadly acting type II-A Acr, which is able to inhibit SpyCas9, SauCas9 as well as SinCas9, although with low potency.

We also observe weak inhibition of SauCas9 with ML3 (AcrIIA12), which was shown to be a specific inhibitor of *Listeria monocytogenes* Cas9 (LmoCas9) while this study was being conducted (25). Because we were unable to test LmoCas9 (due to the difficulty of purifying it intact and active), we were unable to observe strong inhibition activity specific to its host Cas9. Similarly, we were unable to satisfactorily purify *S. agalactiae* Cas9 (SagCas9) to test ML4-ML10 against the Cas9 found in the same genomes in which they were found, leaving the door open for the possibility that they are specific against SagCas9.

The ability to identify potential new Acr candidates directly from protein sequence with AcRanker opens the door for testing many new proteins without the need for laborious screening efforts. Searching within prophages of genomes containing self-targeting CRISPR arrays promises to be particularly effective, as the potential inhibitors for a specific CRISPR system can be quickly ranked to make a short list of candidates to test. We expect that direct Acr prediction methods like AcRanker will continue to reveal many more Acrs distributed across many bacterial species, finding new Acrs with unique properties for yet unforeseen future biotechnology applications.

## DATA AVAILABILITY

A webserver implementation of AcRanker is publicly available at http://acranker.pythonanywhere.com/. The Python code for the webserver implementation is available in the GitHub repository (https://github.com/amina01/AcRanker).

## ACKNOWLEDGEMENTS

We thank Blake McMahon for plasmid cloning and protein purification. We thank Haridha Shivram and Patrick Pausch for providing useful tips throughout the project. We also want to thank Dylan Smock for expressing proteins and Brittney W. Thornton for technical advice.

## AUTHOR CONTRIBUTIONS

Conceptualization, F.A.A.M., K.E.W., J.A.D.; Methodology, A.A., K.E.W., S.E., G.J.K., F.A.A.M.; Software, A.A., F.A.A.M., K.E.W.; Investigation, A.A., K.E.W., S.E., F.A.A.; Biochemical Analysis, S.E., G.J.K., A.T.I.; Data Curation, A.A., K.E.W., S.E.; Writing, A.A., S.E., K.E.W., G.J.K., F.A.A.M.; Funding Acquisition, K.E.W., S.E., G.J.K., J.A.D., F.A.A.M., A.T.I.

