## Supplementary Figures and Tables for "Machine Learning Predicts New Anti-CRISPR Proteins"

*To whom correspondence should be addressed.

[#]Present Address: Department of Computer Science, University of Warwick, Coventry, CV4 7AL, UK

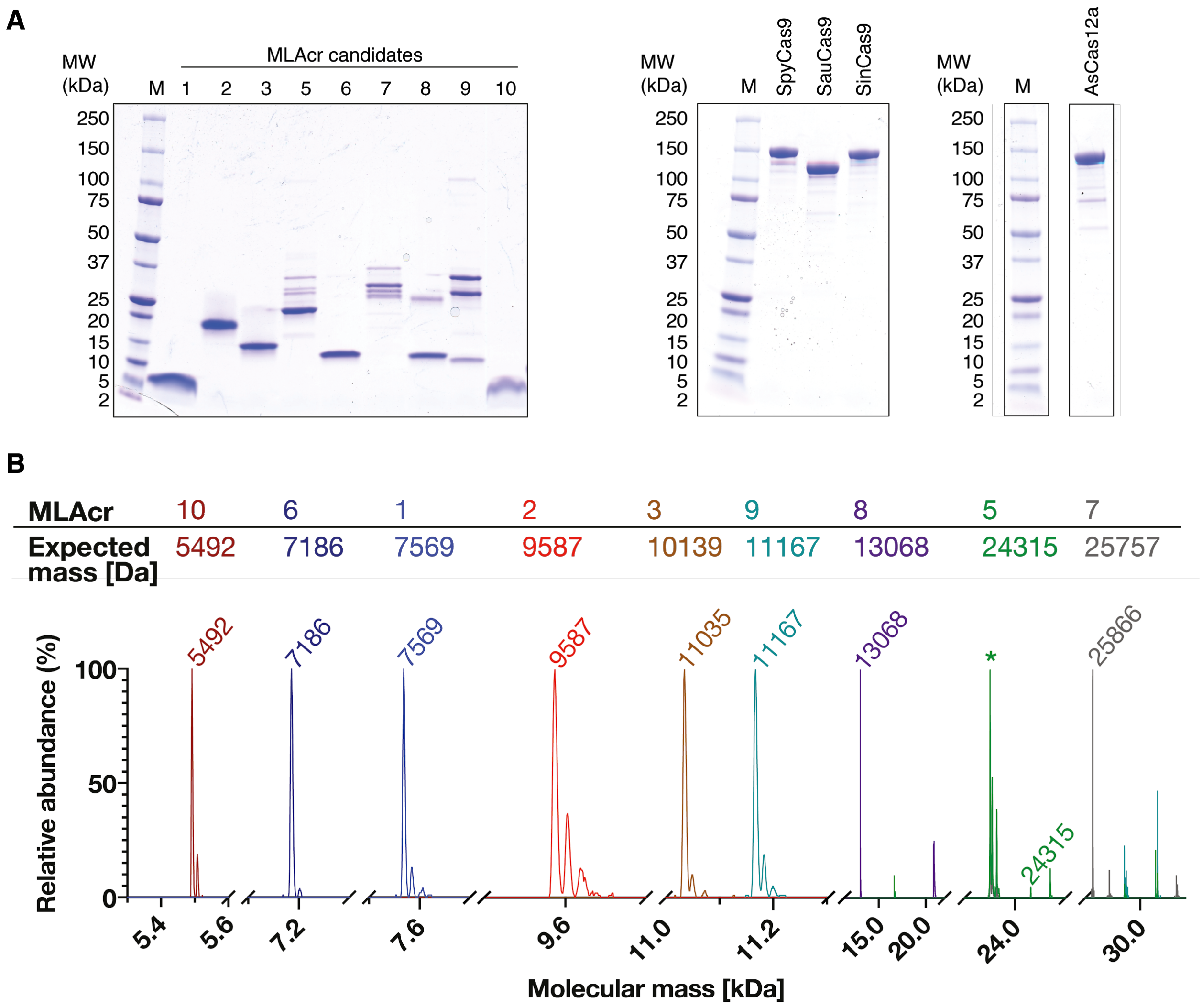

**Figure S1. Purified Acr candidates and Cas effectors used in this study.** (**A**) 4-20% gradient SDS-PAGE showing a size marker (M) and (left to right) purified machine learning Acr candidates, Cas9 effectors and AsCas12a used in this study. (**B**) Mass spectra of each purified Acr candidate used in this study. MLAcr5 contained a significant unidentified contaminant (*) of 23’510 Da in size.

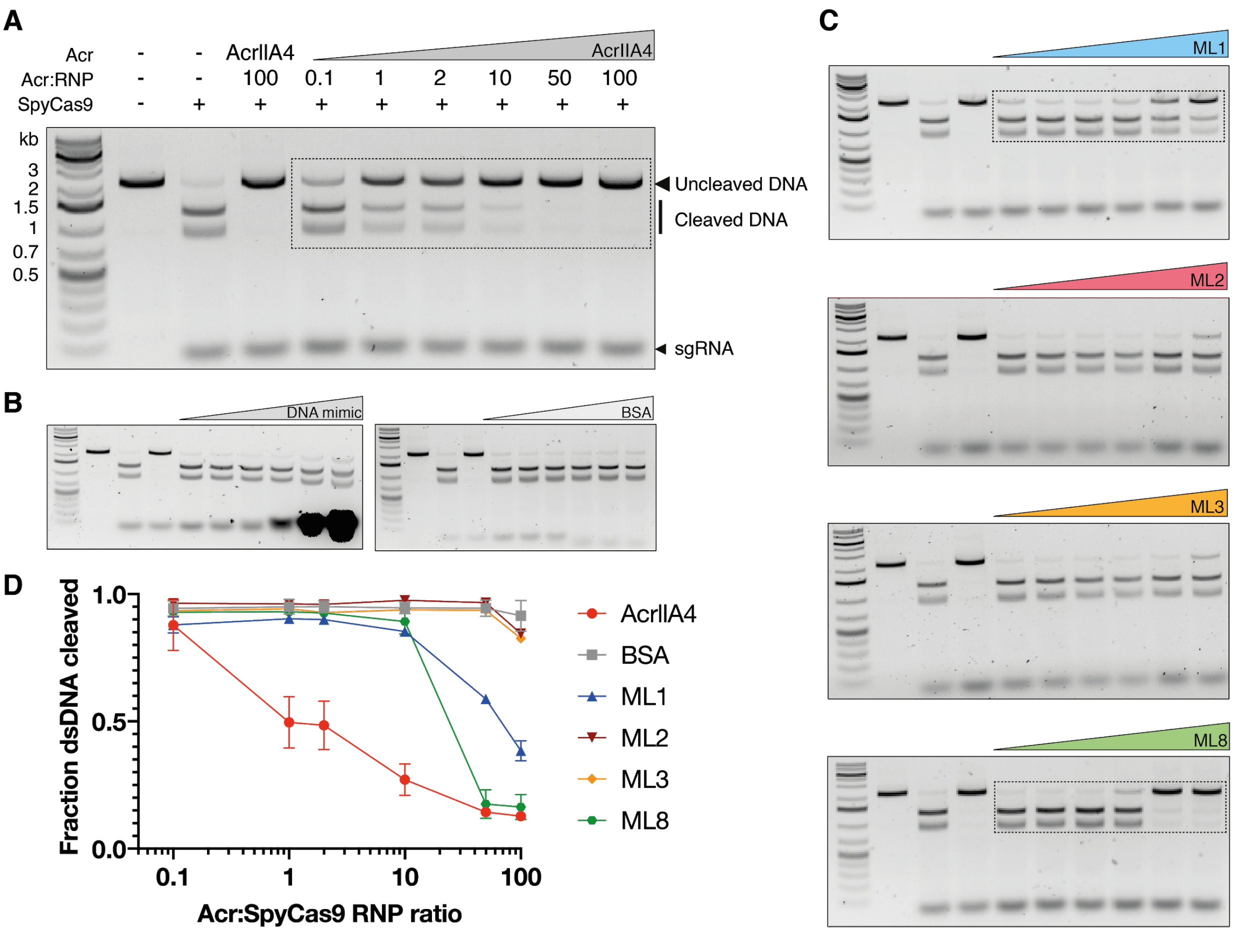

**Figure S2**. **Inhibition of SpyCas9 by newly discovered Acr candidates.** (**A**) *In vitro* cleavage of dsDNA by SpyCas9 in the presence of increasing concentrations of AcrIIA4 (positive control) (**B**) *In vitro* cleavage of dsDNA by SpyCas9 in the presence of increasing concentrations of (left) DNA mimic and (right) BSA (DNA or BSA:RNP 0.1-, 1-, 2- ,10-, 50- and 100-fold excess from left to right). (**C**) *In vitro* cleavage of dsDNA by SpyCas9 in the presence of increasing concentrations of ML1, ML2, ML3 and ML8 (Acr:RNP 0.1-, 1-, 2- ,10-, 50- and 100-fold excess from left to right). (**D**) Quantified band intensities of the *in vitro* cleavage assays. Fraction of dsDNA cleaved (y-axis) is plotted against the Acr to SpyCas9 RNP ratio (x-axis). AcrIIA4, BSA, ML1 and ML8 were run in triplicates.

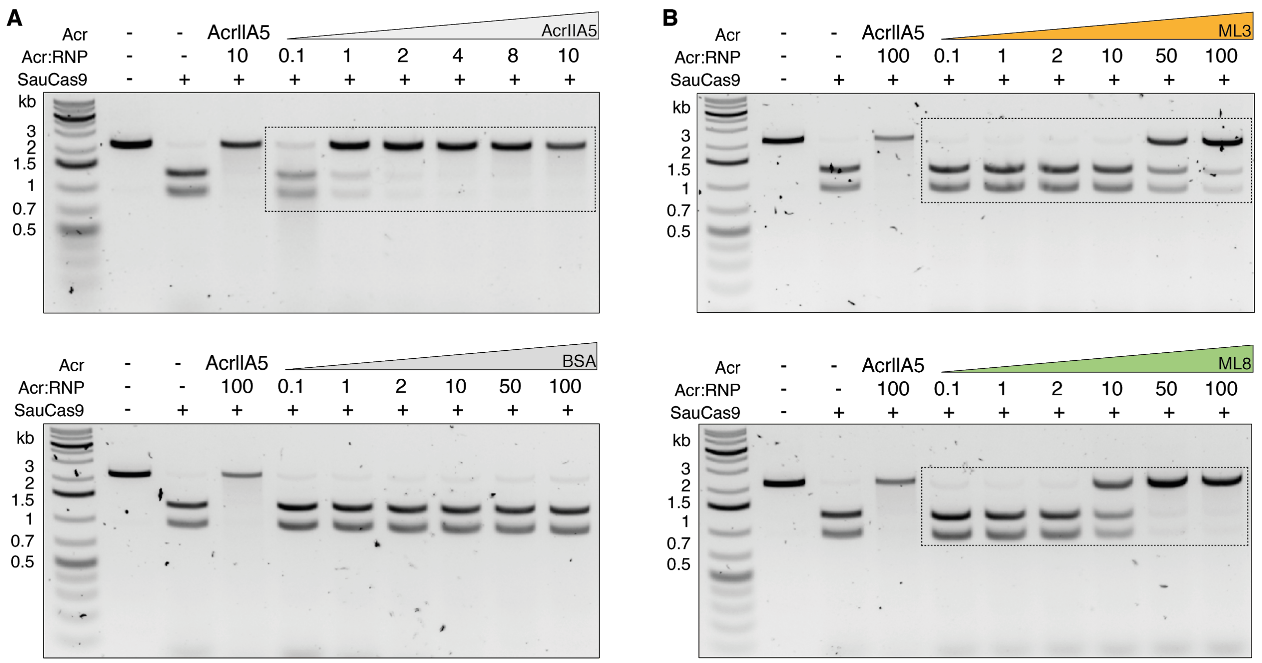

**Figure S3. Inhibition of SauCas9 by newly discovered Acr candidates.** (**A**) *In vitro* cleavage of dsDNA by SauCas9 in the presence of increasing concentrations of the positive control AcrIIA5 (top) or negative control BSA (bottom). (**B**) *In vitro* cleavage of dsDNA by SauCas9 in the presence of increasing concentrations of ML3 (top) and ML8 (bottom).

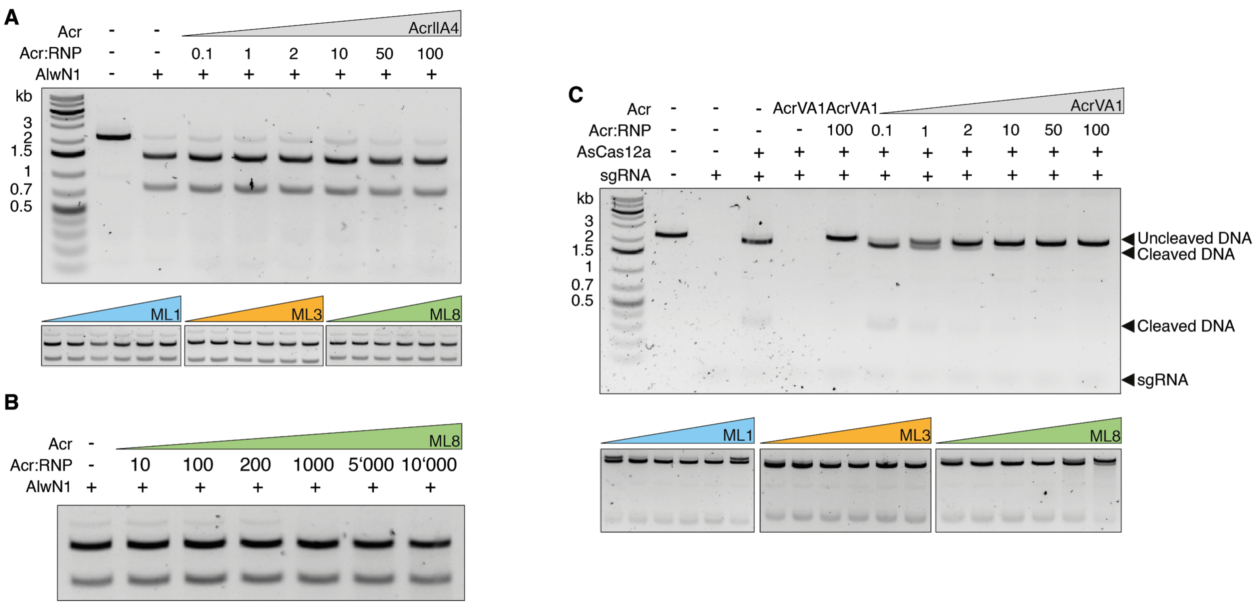

**Figure S4. Control experiments for *in vitro* dsDNA cleavage assay.** (**A**) *In vitro* cleavage of dsDNA by the restriction enzyme AlwN1 in the absence or presence of increasing concentrations of AcrIIA4, ML1, ML3 and ML8 (Acr:AlwN1 0.1-, 1-, 2- ,10-, 50- and 100-fold excess from left to right). (**B**) *In vitro* cleavage of dsDNA by the restriction enzyme AlwN1 in the presence of increasing concentrations of ML8. (**C**) *In vitro* cleavage of dsDNA by AsCas12a in the absence or presence of increasing concentrations of AcrVA1, ML1, ML3 and ML8 (Acr:RNP 0.1-, 1-, 2- ,10-, 50- and 100-fold excess from left to right).

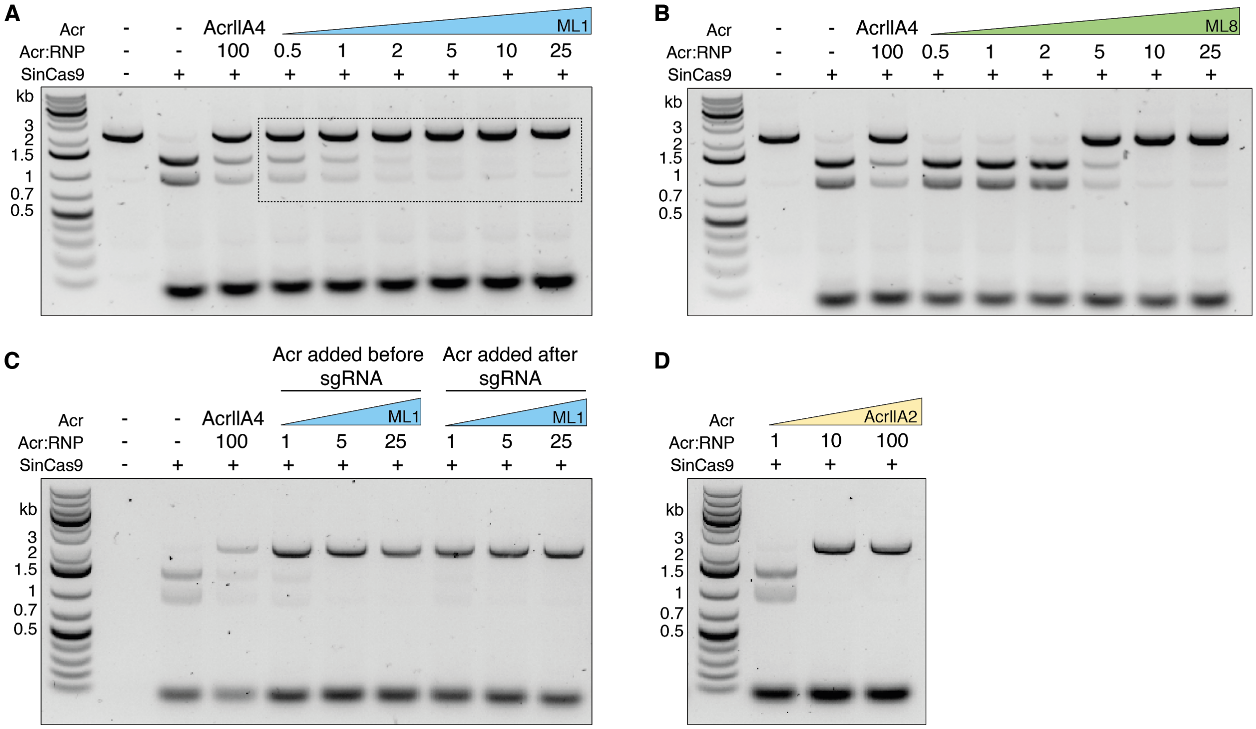

**Figure S5. Inhibition of SinCas9 by ML1, ML8 and AcrIIA2.** (**A**) *In vitro* cleavage of dsDNA by SinCas9 in the absence or presence of increasing concentrations of ML1. (**B**) *In vitro* cleavage of dsDNA by SinCas9 in the absence or presence of increasing concentrations of ML8. (**C**) *In vitro* cleavage assay where ML1 is incubated with SinCas9 before and after the incubation with sgRNA. (**D**) *In vitro* cleavage of dsDNA by SinCas9 in the presence of increasing concentrations of AcrIIA2.

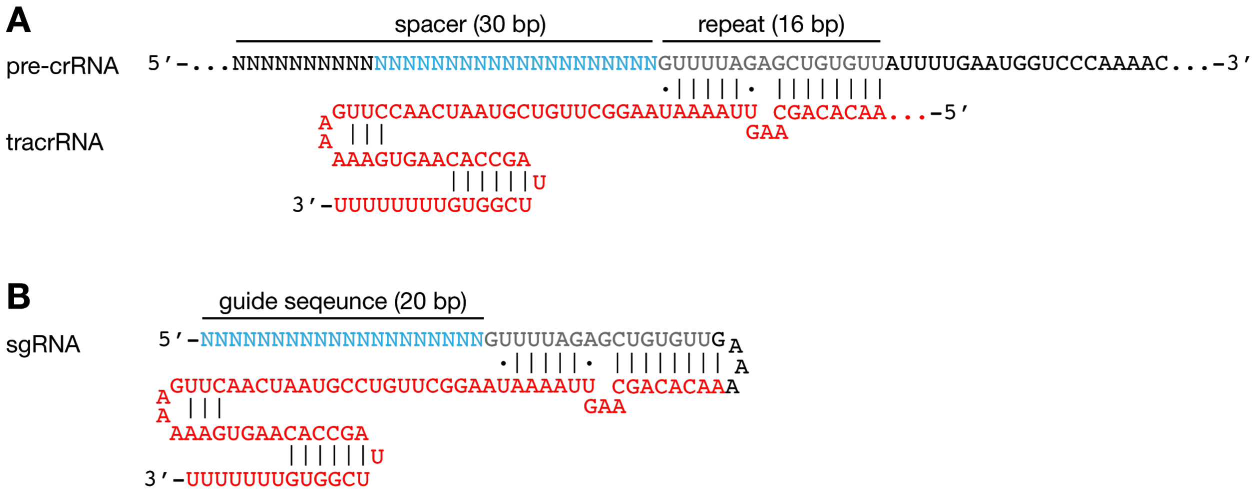

**Figure S6. Design of the SinCas9 single guide RNA.** (**A**) Predicted structure of the SinCas9 pre-crRNA:tracrRNA complex. (**B**) Schematic of the single-guide RNA designed for use in this study. The 20-bp spacer sequence is shown in blue, tracrRNA is shown in red and the direct repeat sequence in grey.

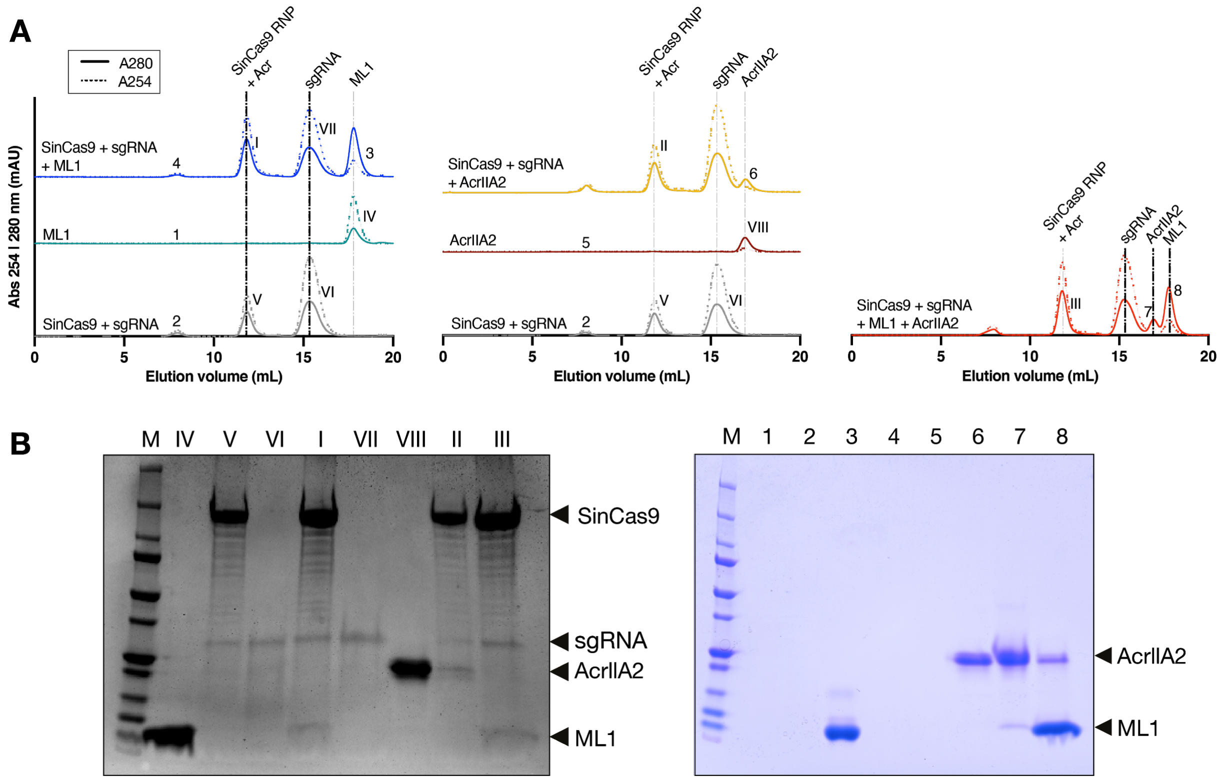

**Figure S7. Competition binding experiment between ML1 and AcrIIA2.** (**A**) Size-exclusion chromatogram of SinCas9-sgRNA in the absence or presence of ML1 (left), AcrIIA2 (middle) or both (right). (**B**) Coomassie-stained polyacrylamide gel illustrating the components of the fractions annotated with (**I**) to (**VI**) and 1 to 8 in panel A.

**Table S1. List of Acrs used for training and cross-validation of the AcRanker model.**

| Anti-CRIPSRdb Name | Acr Family | Protein Accession # | Species | Proteome Size | Ref |
|---|---|---|---|---|---|
| anti_CRISPR0407 | AcrIE1 | YP_007392738.1 | *Pseudomonas* phage JBD5 | 57 | (1) |
| anti_CRISPR0408 | AcrIE3 | YP_950454.1 | *Pseudomonas* phage DMS3 | 52 | (1) |
| anti_CRISPR0409 | AcrIE2 | YP_007392439.1 | *Pseudomonas* phage JBD88a | 54 | (1) |
| anti_CRISPR0410 | AcrIE4 | NP_938238.1 | *Pseudomonas* phage D3112 | 54 | (1) |
| anti_CRISPR0001 | AcrIF1 | YP_007392342.1 | *Pseudomonas* phage JBD30 | 56 | (2) |
| anti_CRISPR0007 | AcrIF2 | YP_002332454.1 | *Pseudomonas* phage MP29 | 51 | (2) |
| anti_CRISPR0003 | AcrIF3 | YP_007392440.1 | *Pseudomonas* phage JBD88a | 54 | (2) |
| anti_CRISPR0002 | AcrIF4 | YP_007392799.1 | *Pseudomonas* phage JBD24 | 57 | (2) |
| anti_CRISPR0005 | AcrIF5 | YP_007392740.1 | *Pseudomonas* phage JBD5 | 57 | (2) |
| anti_CRISPR0008 | AcrIF6 | WP_043884810.1 | *Pseudomonas aeruginosa* | 6095 | (3) |
| anti_CRISPR0011 | AcrIF6 | WP_019933870.1 | *Oceanimonas smirnovii* | 3045 | (3) |
| anti_CRISPR0013 | AcrIF6 | WP_014702809.1 | *Methylophaga frappieri* | 2689 | (3) |
| anti_CRISPR0022 | AcrIF7 | ACD38920.1 | *Pseudomonas aeruginosa strain PACS458 clone fa1376 Pseudomonas aeruginosa* | 57 | (3) |
| anti_CRISPR0034 | AcrIF8 | AFC22483.1 | *Pectobacterium* phage ZF40 | 68 | (3) |
| anti_CRISPR0038 | AcrIF9 | WP_031500045.1 | *Vibrio parahaemolyticus* | 4928 | (3) |
| anti_CRISPR0051 | AcrIF10 | KEK29119.1 | *Shewanella xiamenensis* | 3552 | (3) |
| anti_CRISPR0134 | AcrIIA1 | AEO04364.1 | *Listeria monocytogenes* J0161 | 2952 | (4) |

| | | | | | |
|---|---|---|---|---|---|
| anti_CRISPR0246 | AcrIIA2 | AEO04363.1 | *Listeria monocytogenes* J0161 | 2952 | (4) |
| anti_CRISPR0384 | AcrIIA4 | AEO04689.1 | *Listeria monocytogenes* J0161 | 2952 | (4) |
| anti_CRISPR0433 | AcrIIA5 | D4276_028 | *Streptococcus* phage D4276 | 54 | (5) |

**Table S2. List of Acrs used for independent testing of AcRanker.**

| Acr Family | Protein Accession # | Species | Proteome Size | Ref |
|---|---|---|---|---|
| AcrIE5 | WP_074973300.1 | *Pseudomonas otitidis* strain DSM 17224 | 5731 | (6) |
| AcrIE6 | WP_087937214.1 | *Pseudomonas aeruginosa* strain S708_C14_RS | 6794 | (6) |
| AcrIE7 | WP_087937215.1 | *Pseudomonas aeruginosa* strain S708_C14_RS | 6794 | (6) |
| AcrIE4-F7 | WP_064584002.1 | *Pseudomonas citronellolis* strain SJTE-3 | 6260 | (6) |
| AcrIF11 | WP_038819808.1 | *Pseudomonas aeruginosa* str. C1426 | 5888 | (6) |
| AcrIF11.1 | WP_033936089.1 | *Pseudomonas aeruginosa* strain TRN6649 | 6373 | (6) |
| AcrIF11.2 | EGE18857.1 | *Moraxella catarrhalis* BC8 | 1844 | (6) |
| AcrIF12 | ABR13388.1 | *Pseudomonas aeruginosa* PAGI-5 genomic island sequence | 121 | (6) |
| AcrIF13 | EGE18854.1 | *Moraxella catarrhalis* BC8 | 1843 | (6) |
| AcrIF14 | AKI27193.1 | *Moraxella* phage Mcat5 | 68 | (6) |

**Table S3. Grouping of amino acids based on physiochemical properties.** Groups of amino acids with similar side chains are grouped together to reduce the number of features to test in the machine learning model (7).

| Groups | Amino Acids |
|---|---|
| **1** | A, G, V |
| **2** | I, L, F, P |
| **3** | Y, M, T, S |
| **4** | H, N, Q, W |
| **5** | R, K |
| **6** | D, E |
| **7** | C |

**Table S4. Amino acid sequence and accession numbers of all the Acr candidates.**

| #ML cand. | Accession No. | Sequence |
|---|---|---|
| ML1 | OHX26873.1 | MKNYEVTNEVKNLNTQVETIGQAVDLYKEYGSNTIVWSIDKN EDLIDEVTELVAEYAEKGTVIK |
| ML2 | WP_003731277.1 | MGKTYWYNEGTDTLLTEKEYKELMEREAKALYEEVQEEEKD FESSEKTSFEEFLKTCYENESDFVLSDNEGNKLEEW |
| ML3 | WP_003731276.1 | MSKTMYKNDVIELIKNAKTNNEELLFTSVERNTREAATQYFR CPEKHVSDAGVYYGEDFEFDGFEIFEDDLIYTRSYDKEELN |
| ML4 | WP_000946250.1 | MLRRVNHVKNVLAHGEFAEWIENKIGIHYREANRMMTVAKQI PNVSTLKYLGATAKHVNGVAKRKQNFLSQISLIPTNPQLPHQ TIINTYLYWQP |
| ML5 | WP_001080841.1 | MNRLKELRKEKKLTQEELAGEIGVSKITILRWENGERQIKPDK AKELAKYFNVSVGYLLGYAPNKKIDFQLNLDGTTLHLTKEQFL ALENTSKSIKKIKNTINESVKQEEYIKNASKYYDFEKVSRRLTD RLFEIHTDLIELLMMLDHFPSGELSKSQQEAIFKFYKQLDYFV TDTPASFDYFKKNLESYGYKIYTEGDKIDFD |
| ML6 | WP_000965633.1 | MLYIDEFKEAIDKGYILGGTVAIVRKNGKIFDYVLPHEEVREEE VVTVERVEDVMRELE |
| ML7 | WP_000591144.1 | MIKIYFGKDAALNQAIQSRLDSYQIDYQAFSSKDIDAKTLMEW LFKSTDIFELLSTKMLKYKLNTQITLSQFVRKILKDVNSTLKLPI VVTDEVIYSNMSPDYVTVLLPKEYRKIKRIQLMRKMEQLDEG RLFWKNFELFRKQSELRWFELNELLFADVSDDLGEIKKAKDR FFSYKKNNQVPPNEIIERILKIFLVDREDFFKKSPSDLQNF |
| ML8 | WP_000384271.1 | MDYDNENYLIPKILLQDDFYSSLSAKDILVYAVLKDRQIEALEK GWIDTDGSIYLNFKLIELAKMFSCSRTTMIDVMQRLEEVNLIE RERVDVFYGYSLPYKTYINEV |
| ML9 | WP_000134666.1 | MTEGFTIQLPKVTEKKLLARYDDMLQKAIEKALEDKELYKPIV RMAGLCRWLDVSTTTVVKWQKQGGMPHMVIDGVTLYDKHK VAQWLQQFER |
| ML10 | WP_011058321.1 | MNIEDIERIISEYLIFRSDIDGCAVIDIEDFLKHIRFSYERLK |

**Table S5. Amino acid sequence of all the Cas effectors used in this study.**

| Cas effector | Species | Strain | Sequence |
|---|---|---|---|
| Cas9 | *Streptococcus pyogenes* | | MDKKYSIGLDIGTNSVGWAVITDEYKVPSKKFKV LGNTDRHSIKKNLIGALLFDSGETAEATRLKRTA RRRYTRRKNRICYLQEIFSNEMAKVDDSFFHRL EESFLVEEDKKHERHPIFGNIVDEVAYHEKYPTI YHLRKKLVDSTDKADLRLIYLALAHMIKFRGHFLI EGDLNPDNSDVDKLFIQLVQTYNQLFEENPINAS GVDAKAILSARLSKSRRLENLIAQLPGEKKNGLF GNLIALSLGLTPNFKSNFDLAEDAKLQLSKDTYD DDLDNLLAQIGDQYADLFLAAKNLSDAILLSDILR VNTEITKAPLSASMIKRYDEHHQDLTLLKALVRQ QLPEKYKEIFFDQSKNGYAGYIDGGASQEEFYK FIKPILEKMDGTEELLVKLNREDLLRKQRTFDNG SIPHQIHLGELHAILRRQEDFYPFLKDNREKIEKIL TFRIPYYVGPLARGNSRFAWMTRKSEETITPWN FEEVVDKGASAQSFIERMTNFDKNLPNEKVLPK HSLLYEYFTVYNELTKVKYVTEGMRKPAFLSGE QKKAIVDLLFKTNRKVTVKQLKEDYFKKIECFDS VEISGVEDRFNASLGTYHDLLKIIKDKDFLDNEE NEDILEDIVLTLTLFEDREMIEERLKTYAHLFDDK VMKQLKRRRYTGWGRLSRKLINGIRDKQSGKTI LDFLKSDGFANRNFMQLIHDDSLTFKEDIQKAQV SGQGDSLHEHIANLAGSPAIKKGILQTVKVVDEL VKVMGRHKPENIVIEMARENQTTQKGQKNSRE RMKRIEEGIKELGSQILKEHPVENTQLQNEKLYL YYLQNGRDMYVDQELDINRLSDYDVDHIVPQSF LKDDSIDNKVLTRSDKNRGKSDNVPSEEVVKKM KNYWRQLLNAKLITQRKFDNLTKAERGGLSELD KAGFIKRQLVETRQITKHVAQILDSRMNTKYDEN DKLIREVKVITLKSKLVSDFRKDFQFYKVREINNY HHAHDAYLNAVVGTALIKKYPKLESEFVYGDYK VYDVRKMIAKSEQEIGKATAKYFFYSNIMNFFKT EITLANGEIRKRPLIETNGETGEIVWDKGRDFAT VRKVLSMPQVNIVKKTEVQTGGFSKESILPKRN |

| | | | |
|---|---|---|---|
| | | | SDKLIARKKDWDPKKYGGFDSPTVAYSVLVVAK VEKGKSKKLKSVKELLGITIMERSSFEKNPIDFLE AKGYKEVKKDLIIKLPKYSLFELENGRKRMLASA GELQKGNELALPSKYVNFLYLASHYEKLKGSPE DNEQKQLFVEQHKHYLDEIIEQISEFSKRVILADA NLDKVLSAYNKHRDKPIREQAENIIHLFTLTNLGA PAAFKYFDTTIDRKRYTSTKEVLDATLIHQSITGL YETRIDLSQLGGD |
| Cas9 | *Staphylococcus aureus* | | MGKRNYILGLDIGITSVGYGIIDYETRDVIDAGVR LFKEANVENNEGRRSKRGARRLKRRRRHRIQR VKKLLFDYNLLTDHSELSGINPYEARVKGLSQKL SEEEFSAALLHLAKRRGVHNVNEVEEDTGNELS TKEQISRNSKALEEKYVAELQLERLKKDGEVRG SINRFKTSDYVKEAKQLLKVQKAYHQLDQSFIDT YIDLLETRRTYYEGPGEGSPFGWKDIKEWYEML MGHCTYFPEELRSVKYAYNADLYNALNDLNNLV ITRDENEKLEYYEKFQIIENVFKQKKKPTLKQIAK EILVNEEDIKGYRVTSTGKPEFTNLKVYHDIKDIT ARKEIIENAELLDQIAKILTIYQSSEDIQEELTNLN SELTQEEIEQISNLKGYTGTHNLSLKAINLILDEL WHTNDNQIAIFNRLKLVPKKVDLSQQKEIPTTLV DDFILSPVVKRSFIQSIKVINAIIKKYGLPNDIIIELA REKNSKDAQKMINEMQKRNRQTNERIEEIIRTTG KENAKYLIEKIKLHDMQEGKCLYSLEAIPLEDLLN NPFNYEVDHIIPRSVSFDNSFNNKVLVKQEENSK KGNRTPFQYLSSSDSKISYETFKKHILNLAKGKG RISKTKKEYLLEERDINRFSVQKDFINRNLVDTR YATRGLMNLLRSYFRVNNLDVKVKSINGGFTSF LRRKWKFKKERNKGYKHHAEDALIIANADFIFKE WKKLDKAKKVMENQMFEEKQAESMPEIETEQE YKEIFITPHQIKHIKDFKDYKYSHRVDKKPNRELI NDTLYSTRKDDKGNTLIVNNLNGLYDKDNDKLK KLINKSPEKLLMYHHDPQTYQKLKLIMEQYGDE KNPLYKYYEETGNYLTKYSKKDNGPVIKKIKYYG NKLNAHLDITDDYPNSRNKVVKLSLKPYRFDVYL DNGVYKFVTVKNLDVIKKENYYEVNSKCYEEAK |

| | | | |
|---|---|---|---|
| | | | KLKKISNQAEFIASFYNNDLIKINGELYRVIGVNN<br>DLLNRIEVNMIDITYREYLENMNDKRPPRIIKTIAS<br>KTQSIKKYSTDILGNLYEVKSKKHPQIIKKG |
| Cas9 | *Streptococcus iniae* | UEL-Si1 | MRKPYSIGLDIGTNSVGWAVITDDYKVPSKKMRI<br>QGTTDRTSIKKNLIGALLFDNGETAEATRLKRTT<br>RRRYTRRKYRIKELQKIFSSEMNELDIAFFPRLS<br>ESFLVSDDKEFENHPIFGNLKDEITYHNDYPTIY<br>HLRQTLADRDQKADLRLIYLALAHIIKFRGHFLIE<br>GNLDSENTDVHVLFLNLVNIYNNLFEEDIVETASI<br>DAEKILTSKTSKSRRLENLIAEIPNQKRNMLFGNL<br>VSLALGLTPNFKTNFELLEDAKLQISKDSYEEDL<br>DNLLAQIGDQYADLFIAAKKLSDAILLSDIITVKGA<br>STKAPLSASMVQRYEEHQQDLALLKNLVKKQIP<br>EKYKEIFDNKEKNGYAGYIDGKTSQEEFYKYIKPI<br>LLKLNGTEKLISKLEREDFLRKQRTFDNGSIPHQI<br>HLNELKAIIRRQEKFYPFLKENQKKIEKLFTFKIPY<br>YVGPLANGQSSFAWLKRQSNESITPWNFEEVV<br>DQEASARAFIERMTNFDTYLPEEKVLPKHSPLY<br>EMFMVYNELTKVKYQTEGMKRPVFLSSEDKEEI<br>VNLLFKKDRKVTVKQLKEEYFSKMKCFHTVTILG<br>VEDRFNASLGTYHDLLKIFKDKAFLDDEANQDIL<br>EEIVWTLTLFEDQAMIERRLVKYADVFEKSVLKK<br>LKKRHYTGWGRLSQKLINGIKDKQTGKTILGFLK<br>DDGVANRNFMQLINDSSLDFAKIIKHEQEKTIKN<br>ESLEETIANLAGSPAIKKGILQSIKIVDEIVKIMGQ<br>NPDNIVIEMARENQSTMQGIKNSRQRLRKLEEV<br>HKNTGSKILKEYNVSNTQLQSDRLYLYLLQDGK<br>DMYTGKELDYDNLSQYDIDHIIPQSFIKDNSIDNI<br>VLTTQASNRGKSDNVPNIEIVNKMKSFWYKQLK<br>NGAISQRKFDHLTKAERGALSDFDKAGFIKRQL<br>VETRQITKHVAQILDSRFNSNLTEDSKSNRNVKII<br>TLKSKMVSDFRKDFGFYKLREVNDYHHAQDAY<br>LNAVVGTALLKKYPKLEAEFVYGDYKHYDLAKL<br>MIQPDSSLGKATTRMFFYSNLMNFFKKEIKLADD<br>TIFTRPQIEVNTETGEIVWDKVKDMQTIRKVMSY<br>PQVNIVMKTEVQTGGFSKESILPKGNSDKLIARK |

| | | | |
|---|---|---|---|
| | | | KSWDPKKYGGFDSPIIAYSVLVVAKIAKGKTQKL KTIKELVGIKIMEQDEFEKDPIAFLEKKGYQDIQT SSIIKLPKYSLFELENGRKRLLASAKELQKGNELA LPNKYVKFLYLASHYTKFTGKEEDREKKRSYVE SHLYYFDEIMQIIVEYSNRYILADSNLIKIQNLYKE KDNFSIEEQAINMLNLFTFTDLGAPAAFKFFNGDI DRKRYSSTNEIINSTLIYQSPTGLYETRIDLSKLG GK |
| Cas12a | *Acidaminococcus sp.* | | MTQFEGFTNLYQVSKTLRFELIPQGKTLKHIQEQ GFIEEDKARNDHYKELKPIIDRIYKTYADQCLQLV QLDWENLSAAIDSYRKEKTEETRNALIEEQATYR NAIHDYFIGRTDNLTDAINKRHAEIYKGLFKAELF NGKVLKQLGTVTTTEHENALLRSFDKFTTYFSG FYENRKNVFSAEDISTAIPHRIVQDNFPKFKENC HIFTRLITAVPSLREHFENVKKAIGIFVSTSIEEVF SFPFYNQLLTQTQIDLYNQLLGGISREAGTEKIK GLNEVLNLAIQKNDETAHIIASLPHRFIPLFKQILS DRNTLSFILEEFKSDEEVIQSFCKYKTLLRNENVL ETAEALFNELNSIDLTHIFISHKKLETISSALCDH WDTLRNALYERRISELTGKITKSAKEKVQRSLKH EDINLQEIISAAGKELSEAFKQKTSEILSHAHAAL DQPLPTTLKKQEEKEILKSQLDSLLGLYHLLDWF AVDESNEVDPEFSARLTGIKLEMEPSLSFYNKA RNYATKKPYSVEKFKLNFQMPTLASGWDVNKE KNNGAILFVKNGLYYLGIMPKQKGRYKALSFEPT EKTSEGFDKMYYDYFPDAAKMIPKCSTQLKAVT AHFQTHTTPILLSNNFIEPLEITKEIYDLNNPEKEP KKFQTAYAKKTGDQKGYREALCKWIDFTRDFLS KYTKTTSIDLSSLRPSSQYKDLGEYYAELNPLLY HISFQRIAEKEIMDAVETGKLYLFQIYNKDFAKGH HGKPNLHTLYWTGLFSPENLAKTSIKLNGQAEL FYRPKSRMKRMAHRLGEKMLNKKLKDQKTPIP DTLYQELYDYVNHRLSHDLSDEARALLPNVITKE VSHEIIKDRRFTSDKFFFHVPITLNYQAANSPSKF NQRVNAYLKEHPETPIIGIDRGERNLIYITVIDSTG KILEQRSLNTIQQFDYQKKLDNREKERVAARQA |

| | | | |
|---|---|---|---|
| | | | WSVVGTIKDLKQGYLSQVIHEIVDLMIHYQAVVV LENLNFGFKSKRTGIAEKAVYQQFEKMLIDKLNC LVLKDYPAEKVGGVLNPYQLTDQFTSFAKMGTQ SGFLFYVPAPYTSKIDPLTGFVDPFVWKTIKNHE SRKHFLEGFDFLHYDVKTGDFILHFKMNRNLSF QRGLPGFMPAWDIVFEKNETQFDAKGTPFIAGK RIVPVIENHRFTGRYRDLYPANELIALLEEKGIVF RDGSNILPKLLENDDSHAIDTMVALIRSVLQMRN SNAATGEDYINSPVRDLNGVCFDSRFQNPEWP MDADANGAYHIALKGQLLLNHLKESKDLKLQNGI SNQDWLAYIQELRN |

**Table S6. sgRNAs used for the *in vitro* cleavage assay.**

| Cas effector | Species | Strain | sgRNA sequence* | Main Text Fig. | Suppl. Fig. |
|---|---|---|---|---|---|
| Cas9 | *Streptococcus pyogenes* | | **ATACGGGAGGGCTTACCATC**GTTTTAGAGCTATGCTGTTTTGGAAACAAAACAGCATAGCAAGTTAAAATAAGGCTAGTCCGTTATCAACTTGAAAAAGTGGCACCGAGTCGGTGCTTTTTTT | 3A-B | 1A, 2A-D |
| Cas9 | *Staphylococcus aureus* | | **TATCGTAGTTATCTACACGACG**GTTTTAGTACTCTGGAAACAGAATCTACTAAAACAAGGCAAAATGCCGTGTTTATCTCGTCAACTTGTTGGCGAGATTTTT | 3C-D | 1A, 3A-B |
| Cas9 | *Streptococcus iniae* | UEL-Si1 | **ATACGGGAGGGCTTACCATC**GTTTTAGAGCTGTGTTGAAAAACACAGCAAGTTAAAATAAGGCTTGTCCGTAATCAACTTGAAAAAGTGAACACCGATTCGGTGTTTTTTT | 4A-B | 5A-B |
| Cas12a | *Acidaminococcus sp.* | | AAUUUCUACUCUUGUAGAU**AAAGUGCUCAUCAUUGGAAAACGU** | - | 4C |

* Spacer sequences are shown in bold

**Table S7. DNA target used for the *in vitro* cleavage assay.**

<table>
<tr><th>Cas effector</th><th>Species</th><th>DNA target sequence*</th><th>Main Text Fig.</th><th>Suppl. Fig.</th></tr>
<tr><td>Cas9</td><td>Streptococcus pyogenes</td><td rowspan="4">AATAATGGTTTCTTAGACGTCAGGTGGCACTTTTC<br>GGGGAAATGTGCGCGGAACCCCTATTTGTTTATTT<br>TTCTAAATACATTCAAATATGTATCCGCTCATGAGA<br>CAATAACCCTGATAAATGCTTCAATAATATTGAAAA<br>AGGAAGAGTATGAGTATTCAACATTTCCGTGTCGC<br>CCTTATTCCCTTTTTTGCGGCATTTTGCCTTCCTGT<br>TTTTGCTCACCCAGAAACGCTGGTGAAAGTAAAAG<br>ATGCTGAAGATCAGTTGGGTGCACGAGTGGGTTA<br>CATCGAACTGGATCTCAACAGCGGTAAGATCCTTG<br>AGAGTTTTCGCCCCGAAGAACGTTTTCCAATGATG<br>AGCACTTTTAAAGTTCTGCTATGTGGCGCGGTATT<br>ATCCCGTATTGACGCCGGGCAAGAGCAACTCGGT<br>CGCCGCATACACTATTCTCAGAATGACTTGGTTGA<br>GTACTCACCAGTCACAGAAAAGCATCTTACGGATG<br>GCATGACAGTAAGAGAATTATGCAGTGCTGCCATA<br>ACCATGAGTGATAACACTGCGGCCAACTTACTTCT<br>GACAACGATCGGAGGACCGAAGGAGCTAACCGCT<br>TTTTTGCACAACATGGGGGATCATGTAACTCGCCT<br>TGATCGTTGGGAACCGGAGCTGAATGAAGCCATA<br>CCAAACGACGAGCGTGACACCACGATGCCTGTAG<br>CAATGGCAACAACGTTGCGCAAACTATTAACTGGC<br>GAACTACTTACTCTAGCTTCCCGGCAACAATTAATA<br>GACTGGATGGAGGCGGATAAAGTTGCAGGACCAC<br>TTCTGCGCTCGGCCCTTCCGGCTGGCTGGTTTATT<br>GCTGATAAATCTGGAGCCGGTGAGCGTGGGTCTC<br>GCGGTATCATTGCAGCACTGGGGCCAGATGGTAA<br>GCCCTCCCGTATCGTAGTTATCTACACGACGGGG<br>AGTCAGGCAACTATGGATGAACGAAATAGACAGAT<br>CGCTGAGATAGGTGCCTCACTGATTAAGCATTGGT<br>AACTGTCAGACCAAGTTTACTCATATATACTTTAGA<br>TTGATTTAAAACTTCATTTTTAATTTAAAAGGATCTA<br>GGTGAAGATCCTTTTTGATAATCTCATGACCAAAAT</td><td>3A-B</td><td>1A, 2A-D</td></tr>
<tr><td>Cas9</td><td>Staphylococcus aureus</td><td>3C-D</td><td>1A, 3A-B</td></tr>
<tr><td>Cas9</td><td>Streptococcus iniae UEL-Si1</td><td>4A-B</td><td>5A-B</td></tr>
<tr><td>Cas12a</td><td>Acidaminococcus sp.</td><td>-</td><td>4C</td></tr>
</table>

| | | | | |
|---|---|---|---|---|
| | | CCCTTAACGTGAGTTTTCGTTCCACTGAGCGTCAG<br>ACCCCGTAGAAAAGATCAAAGGATCTTCTTGAGAT<br>CCTTTTTTTCTGCGCGTAATCTGCTGCTTGCAAACA<br>AAAAAACCACCGCTACCAGCGGTGGTTTGTTTGCC<br>GGATCAAGAGCTACCAACTCTTTTTCCGAAGGTAA<br>CTGGCTTCAGCAGAGCGCAGATACCAAATACTGTC<br>CTTCTAGTGTAGCCGTAGTTAGGCCACCACTTCAA<br>GAACTCTGTAGCACCGCCTACATACCTCGCTCTGC<br>TAATCCTGTTACCAGTGGCTGCTGCCAGTGGCGAT<br>AAGTCGTGTCTTACCGGGTTGGACTCAAGACGATA<br>GTTACCGGATAAGGCGCAGCGGTCGGGCTGAACG<br>GGGGGTTCGTGCACACAGCCCAGCTTGGAGCGAA<br>CGACCTACACCGAACTGAGATACCTACAGCGTGA<br>GCTATGAGAAAGCGCCACGCTTCCCGAAGGGAGA<br>AAGGCGGACAGGTATCCGGTAAGCGGCAGGGTC<br>GGAACAGGAGAGCGCACGAGGGAGCTTCCAGGG<br>GGAAACGCCTGGTATCTTTATAGTCCTGTCGGGTT<br>TCGCCACCTCTGACTTGAGCGTCGATTTTTGTGAT<br>GCTCGTCAGGGGGGCGGAGCCTGTGGAAAAACG<br>CCAGCAACGCGGCCTTTTTACGGTTCCTGGCCTTT<br>TGCTGGCCTTTTGCTCACATGTTCTTTCCTGCGTTA<br>TCCCCTGATTCTGTGGATAACCGTATTACCGCAGA<br>GTTTGTAGAAACGCAAAAAGGCCATCCGTCAGGAT<br>GGCCTTCTGCTTAATTTGATGCCTGGCAGTTTATG<br>GCGGGCGTCCTGCCCGCCACCCTCCGGGCCGTT<br>GCTTCGCAACGTTCAAATCCGCTCCCGGCGGTTG<br>AGAAGAGAAAAGAAAACCGCCGATCCTGTCCACC<br>GCATTACTGCAAGGTAGTGGACAAGACCGGCGGT<br>CTTAAGTTTTTTGGCTGAAATGCCTGGCAGTTCCC<br>TACTCTCGCATGGGGCTCGCGGTTAACTGATTATT<br>TTATTTATCTAGGCTACTTACGAACG | | |
| | | **DNA mimic****<br>GCTGACAATGATACGAACGAGACACACGCTCACG<br>ACTCAG | - | 2B |

* Target sequences are shown in blue (*Streptococcus pyogenes, Streptococcus iniae*), yellow (*Acidaminococcus sp.*) or bold (*Staphylococcus aureus*); **DNA mimic used for control experiments

**Table S8. Independent testing set validation results**. Ten proteomes containing non-redundant (<40% sequence identity) Acrs from bacterial and phage sources were ranked using AcRanker. Bacterial proteomes that had Acrs within PHASTER-predicted prophages were also tested with a subset of the proteome containing only the prophage proteins.

| | | Complete Proteome | | Prophage Subset | |
|---|---|---|---|---|---|
| **Anti-CRISPR ID** | **Anti-CRISPR family** | **Proteome Size** | **AcRanker rank** | **Proteome Size** | **AcRanker rank** |
| WP_064584002.1 | AcrIE4-F7 | 6260 | 68 | 111 | 4 |
| WP_038819808.1 | AcrIF11 | 5888 | 138 | 64 | 3 |
| WP_033936089.1 | AcrIF11.1 | 6373 | 38 | 92 | **1** |
| EGE18857.1 | AcrIF11.2 | 1844 | 412 | 59 | 30 |
| AKI27193.1 | AcrIF14 | 68 | 14 | 68 | 14 |
| WP_074973300.1 | AcrIE5 | 5731 | 10 | - | - |
| WP_087937214.1 | AcrIE6 | 6794 | 80 | - | - |
| WP_087937215.1 | AcrIE7 | 6794 | 742 | - | - |
| ABR13388.1 | AcrIF12 | 121 | 7 | - | - |
| EGE18854.1 | AcrIF13 | 1843 | 187 | - | - |

**Table S9. List of expected lethal self-targeting *Streptococcus* genomes obtained with Self-Target Spacer Searcher (STSS).** Searching *Streptococcus* assemblies from NCBI with STSS returned 385 cases of self-targeting derived from type II-A arrays representing 241 individual genomes. Of those genomes, 20 contained at least one spacer with the characteristic NRG 3′ PAM for SpyCas9, shown in the table below. Only *Streptococcus iniae* strain UEL-Si1 contains a previously discovered anti-CRISPR (AcrIIA3). Also shown in the table are the self-targeting spacers for *Listeria monocytogenes* strain R2-502, which was also ranked with AcRanker.

| Target Accession# | Locus Accession# | Species/Strain | Self-Targeting Spacer Sequence(s) | 3′ PAM Region | Anti-CRISPRs Present |
|---|---|---|---|---|---|
| NZ_MNAC01000031.1 | NZ_MNAC01000010.1 | *Streptococcus iniae* strain UEL-Si1 | TTGATAAGTATAATTTCCTGTCTTTGTTTT | AGGAGTTTT | AcrIIA3 (WP_071127625.1) |
| NZ_MNAC01000046.1 | | | TAAGGAATTTGAAGCAATACGTCTTAATTT | AGCAATGAC | |
| NZ_MNAC01000023.1 | | | CAAAAAAGTTCGGTAACTTACGGTAACTTA | CGGTAACTT | |
| | | | TCTAAAAAATCAAAAGTTACCGTGTTACCG | TAGTTTTGA | |
| | | | AATATGACTTTTGGGAAATTAACTAATCAA | TGGCTGAAA | |
| | | | TTTTTGAGTGTACTGATGTTGCTTTTGAGC | TGGCCACTT | |
| NZ_MNAC01000021.1 | | | ATAATCAATCACATTAATGCTGACATCAAC | TGGAGCAGA | |
| | | | GAGTTTAATTAAGTGACATAATATCTTCAT | CGGTTATAG | |
| NZ_JRLL01000002.1 | NZ_JRLL01000058.1 | *Streptococcus pyogenes* SS1447 | TCGTCAGATTTGTCAGTATAGTAATCATCA | CGATATAAA | None |
| NZ_JRLL01000072.1 | | | CTATATTGTTGAGCTGTGGGCTTTGCATAA | AGGTTTAAA | |

| | | | | | |
|---|---|---|---|---|---|
| NZ_JRLL01000026.1 | | | GTAATAATAGCATTGCCTGTTCTATCCTGT | CGGTAGAAC | |
| NZ_CQAV01000003.1 | NZ_CQAV01000001.1 | *Streptococcus agalactiae* strain DE-NI-032 | TATTTGATAGCGGTAACGGGTCATATACAA | AGGCATCTA | None |
| | | | TGGTGGTATTTATAATGTACGAGCAAATCG | AGGCGCTCC | |
| | | | ACCTTGCTCCGATGACACCATCGCGAACCT | TGGTCTAAT | |
| NZ_CP010449.1 | NZ_CP010449.1 | *Streptococcus pyogenes* strain NGAS322 | ATCGTAAGGCAACAGATTATCGTAAGATCT | AGGTGTATA | None |
| NZ_ALQN01000014.1 | NZ_ALQN01000018.1 | Streptococcus agalactiae CCUG 37430 | ATTTGCAACTTTCTCAAGTGTTGCGAGAGA | TGGAGAATT | None |
| NZ_ALQN01000018.1 | | | GCAAGCACTAAATGAAGCTACTAGACTTAA | AGGTCGCAG | |
| | | | TAATGACATGTGGATTGATATCTCAGAGAA | CGGCGATTA | |
| | | | TGTCATTGTTAAAATCATTTGCATATTTTT | TGGATATAA | |
| | | | TACTTGACGAATTGAAGATGACGGAATTTA | TTGCTCCAC | |
| NZ_CPVL01000019.1 | NZ_CPVL01000003.1 | *Streptococcus agalactiae* strain DE-NI-007 | AAGGCACGCGCAAGATGAATTCATTTCTAA | TGGCTACAC | None |
| | | | TGATGTTCTTTATCAAACATTCTAAATACT | TGGAAGCCC | |
| | | | GAGCCTTGCTTGAGTTTGTGGAGCTTTATA | GGGATGGAA | |
| | | | GTATAATTTAGTTAAGCTTAAATTTAACCA | AGGAGACGT | |

| | | | | | |
|---|---|---|---|---|---|
| NZ_ANCM01000101.1 | NZ_ANCM01000101.1 | *Streptococcus agalactiae* FSL S3-586 | GAAAAAGGCGATGTAGCTTAGAAAGGAGAA | GGGATGGAA | None |
| NZ_ANCM01000006.1 | | | GAAAAAGGCGATGTAGCTTAGAAAGGAGAA | CACCATGAA | |
| NZ_ANCM01000028.1 | | | TACGAAAAGGTTGTGATAAAAGCCATATCA | TCGAGTTTG | |
| NZ_ALTM01000012.1 | NZ_ALTM01000016.1 | *Streptococcus agalactiae* GB00548 | AACAACTTTCTTACAAAAGGTTCTAGTTTTCTT | TCGCAAAAC | |
| NZ_ALTM01000013.1 | | | ACGCTCTGAGGCAGATGAGGAACAGGCGCA | TAGGCACCC | |
| NZ_ALUZ01000056.1 | NZ_ALUZ01000054.1 | *Streptococcus agalactiae* GB00984 | TGAAAACAAGCGCAAAGCTGTCAGAAAACA | CGGAACTAA | None |
| | | | TACTTGACGAATTGAAGATGACGGAATTTA | TGGCTCCAC | |
| NZ_ALRF01000019.1 | NZ_ALRF01000066.1 | *Streptococcus agalactiae* BSU188 | GAAACTTCGATTAGTTTGCGTACTCGCTCA | CGGCAAAAC | None |
| NZ_ANEM01000019.1 | NZ_ANEM01000012.1 | *Streptococcus agalactiae* MRI Z1-022 | TTGCTGCTAGACCCAAACAGTTTATTTTTAG | GGCCAAAAA | None |
| NZ_ANEM01000074.1 | | | TATTTCATCATAGAAAATCCTGCTAGTGGT | CGGTTATGG | |
| NZ_CQEL01000006.1 | NZ_CQEL01000002.1 | *Streptococcus agalactiae* strain DK-NI-014 | ACACCTAGTTTCAAGTTTTTAGCAGATTTTTT | GGTTACATT | None |
| NZ_CQEL01000008.1 | | | ACGCTCTGAGGCAGATGAGGAACAGGCGCA | TAGGCACCC | |
| NZ_MAWX01000026.1 | NZ_MAWX01000055.1 | | ATTGACTGTTTACGATTTCCTTCCACCGTT | GGGTACAAA | None |

| | | | | | |
|---|---|---|---|---|---|
| | | *Streptococcus agalactiae* strain DK-PW-096 | TGATGAGATTTTTAAAAGACTCACTGATAT | AGGATTGAC | |
| | | | CGCTTAGATGAAGTACAGATTGTAACAAGT | TCGGAAGTA | |
| NZ_CTJD01000013.1 | NZ_CTJD01000001.1 | *Streptococcus agalactiae* strain GB-NI-015 | TGAAAACAAGCGCAAAGCTGTCAGAAAACA | CGGAACTAA | None |
| | | | TACTTGACGAATTGAAGATGACGGAATTTA | TGGCTCCAC | |
| NZ_CPZS01000003.1 | NZ_CPZS01000001.1 | *Streptococcus agalactiae* strain IT-NI-009 | TATTTGATAGCGGTAACGGGTCATATACAA | AGGCATCTA | None |
| | | | ACCTTGCTCCGATGACACCATCGCGAACCT | TGGTCTAAT | |
| NZ_CPVQ01000026.1 | NZ_CPVQ01000002.1 | *Streptococcus agalactiae* strain RBH12 | AACACAGCTTCCTCGAAAGGGATATATCTA | CGGACAACT | None |
| NDGB01000049.1 | NDGB01000023.1 | *Streptococcus agalactiae* strain ST 618 | ATTAAGTTGCTTAGTGCTTTCATAATCATC | TGGAATAAC | None |
| NDGB01000030.1 | | | ATTAAGTTGCTTAGTGCTTTCATAATCATC | TGGAATAAC | |
| NZ_KQ969340.1 | NZ_KQ969342.1 | *Streptococcus oralis* strain DD14 | TTCCATTTCTGATTTGATTCAACAGCAGCA | GGAAATCCT | None |
| | | | TACAGCGGATACAACCCCACCAATAGCCTC | AGGAATTGC | |
| NZ_KQ961462.1 | NZ_KQ961485.1 | *Streptococcus pasteurianus* strain GED7275A | TTTATTCGGCATCGGCTGGTGTTATGGACT | TGGCTGCGG | None |

| | | | | | |
|---|---|---|---|---|---|
| NZ_AWTL01000007.1 | NZ_AWTL01000011.1 | *Streptococcus pyogenes* GA03805 | TAGAGTAAACCGAATCTTTGCCATCTCTGG | CAGTTTGAC | None |
| NZ_LRGN01000012.1 | NZ_LRGN01000001.1 | *Streptococcus pyogenes* strain SST2091-1 | TAGAGTAAACCGAATCTTTGCCATCTCTGG | CAGTTTGAC | None |
| LRGT01000330.1 | LRGT01000062.1 | *Streptococcus pyogenes* strain SST2097-1 | TGGTCTAACTGCGTCTGGTCTGTGAATGA | TAGGTACAA | None |
| NC_021838.1 | NC_021838.1 | *Listeria monocytogenes* strain R2-502 | GGTAAAACAAGCATCGGCGAAGCAGTAACA | TGGCTTCTT | AcrIIA3 (WP_023553812.1), AcrIIA2 (WP_023553814.1), AcrIIA1 (WP_003722518.1), AcrIIA1 (WP_012581438.1) |
| | | | GGTAAAACAAGCATCGGCGAAGCAGTAACA | TGGCTACTC | |
| | | | TAGGTTTAGGGAGTAAATTAGCTCCTTTGG | CAGCTGGGT | |
| | | | TAACTTTAGATACTGCTAAAGAATTAGCAA | TGGTGCAAA | |
| | | | TTGGGCAAAATGACCGTAATAAATCCATTC | CGGTTCATC | |
| | | | TAGGTTTAGGGAGTAAATTAGCTCCTTTGG | CGGCTGGAT | |

**Table S10. Top Acr gene candidates within each genome ranked by AcRanker.** The proteins found within the prophages of 20 *Streptococcus* genomes were ranked using AcRanker; up to the top 10 highest ranking genes are listed in ascending order. Known Acr genes and the 10 genes synthesized for biochemical testing are indicated in the rightmost column. Genomes with fewer than 10 listed have very few annotated proteins found within predicted prophages.

| Organism | Source Contig | Protein | Rank | Candidate # or Acr |
|---|---|---|---|---|
| *Streptococcus iniae* strain UEL-Si1 | NZ_MNAC01000021.1 | WP_071127623.1 | 1 | ML1 |
| | NZ_MNAC01000023.1 | WP_071127667.1 | 2 | |
| | | WP_071127683.1 | 3 | |
| | NZ_MNAC01000021.1 | WP_071127625.1 | 4 | AcrIIA3 |
| | | WP_071127624.1 | 5 | |
| | NZ_MNAC01000023.1 | WP_071127689.1 | 6 | |
| | NZ_MNAC01000021.1 | WP_071127610.1 | 7 | |
| | NZ_MNAC01000023.1 | WP_071127674.1 | 8 | |
| | | WP_071127671.1 | 9 | |
| | | WP_071127685.1 | 10 | |
| *Streptococcus pyogenes* strain SS1447 | NZ_JRLL01000026.1 | WP_029713970.1 | 1 | |
| | | WP_003057301.1 | 2 | |
| | | WP_032460878.1 | 3 | |
| | NZ_JRLL01000072.1 | WP_002986828.1 | 4 | |
| | NZ_JRLL01000026.1 | WP_012678849.1 | 5 | |
| | | WP_032460877.1 | 6 | |
| | | WP_032460876.1 | 7 | |
| | NZ_JRLL01000072.1 | WP_002994100.1 | 8 | |
| | NZ_JRLL01000026.1 | WP_032460879.1 | 9 | |
| | | WP_050491258.1 | 10 | |
| *Streptococcus agalactiae* strain DE-NI-032 | NZ_CQAV01000003.1 | WP_017827941.1 | 1 | |
| | | WP_050201842.1 | 2 | |
| | | WP_050305756.1 | 3 | |
| | | WP_000138374.1 | 4 | |
| | | WP_000258802.1 | 5 | |
| | | WP_025196275.1 | 6 | |
| | | WP_050198474.1 | 7 | |
| | | WP_000695684.1 | 8 | |

| | | | | |
|---|---|---|---|---|
| | | WP_047200442.1 | 9 | |
| | | WP_001921522.1 | 10 | |
| *Streptococcus pyogenes* strain NGAS322 | NZ_CP010449.1 | WP_002984315.1 | 1 | |
| | | WP_011054546.1 | 2 | |
| | | WP_053308468.1 | 3 | |
| | | WP_011017567.1 | 4 | |
| | | WP_032461328.1 | 5 | |
| | | WP_011017568.1 | 6 | |
| | | WP_011017568.1 | 7 | |
| | | WP_011889039.1 | 8 | |
| | | WP_011284878.1 | 9 | |
| | | WP_011017985.1 | 10 | |
| *Streptococcus agalactiae* strain CCUG 37430 | NZ_ALQN01000018.1 | WP_000649300.1 | 1 | |
| | | WP_079261174.1 | 2 | |
| | | WP_000660740.1 | 3 | |
| | | WP_000076700.1 | 4 | |
| | | WP_000033707.1 | 5 | |
| | | WP_000343312.1 | 6 | |
| | | WP_000323860.1 | 7 | |
| | | WP_000879943.1 | 8 | |
| | | WP_001198776.1 | 9 | |
| | | WP_000229419.1 | 10 | |
| *Streptococcus agalactiae* strain DE-NI-007 | NZ_CPVL01000019.1 | WP_000359663.1 | 1 | |
| | | WP_000141918.1 | 2 | |
| | | WP_000205000.1 | 3 | |
| | | WP_000130289.1 | 4 | |
| | | WP_000946250.1 | 5 | ML4 |
| | | WP_001080841.1 | 6 | ML5 |
| | | WP_000206026.1 | 7 | |
| | | WP_000877491.1 | 8 | |
| | | WP_000051914.1 | 9 | |
| | | WP_000983174.1 | 10 | |
| *Streptococcus agalactiae* FSL S3-586 | NZ_ANCM01000028.1 | WP_017643458.1 | 1 | |
| | | WP_000134940.1 | 2 | |
| | | WP_001875290.1 | 3 | |
| | | WP_000789102.1 | 4 | |

| | | | | |
|---|---|---|---|---|
| | NZ_ANCM01000101.1 | WP_003051787.1 | 5 | |
| | NZ_ANCM01000028.1 | WP_000032136.1 | 6 | |
| | | WP_000342242.1 | 7 | |
| | NZ_ANCM01000006.1 | WP_000686776.1 | 8 | |
| | NZ_ANCM01000101.1 | WP_000988928.1 | 9 | |
| | NZ_ANCM01000028.1 | WP_017643459.1 | 10 | |
| *Streptococcus agalactiae* strain GB00548 | NZ_ALTM01000002.1 | WP_000793595.1 | 1 | |
| | | WP_000384271.1 | 2 | ML8 |
| | | WP_000134666.1 | 3 | ML9 |
| | | WP_000035940.1 | 4 | |
| | | WP_017645945.1 | 5 | |
| | | WP_000022172.1 | 6 | |
| *Streptococcus agalactiae* strain GB00984 | NZ_ALUZ01000056.1 | WP_000660738.1 | 1 | |
| | | WP_017827941.1 | 2 | |
| | | WP_000138374.1 | 3 | |
| | | WP_000614971.1 | 4 | |
| | | WP_000258802.1 | 5 | |
| | | WP_000763911.1 | 6 | |
| | | WP_000239531.1 | 7 | |
| | | WP_000159371.1 | 8 | |
| | | WP_000695684.1 | 9 | |
| | | WP_000698337.1 | 10 | |
| *Streptococcus agalactiae* strain BSU188 | NZ_ALRF01000068.1 | WP_000965633.1 | 1 | ML6 |
| | | WP_000274022.1 | 2 | |
| | | WP_000076712.1 | 3 | |
| | | WP_001183891.1 | 4 | |
| | | WP_000763914.1 | 5 | |
| | | WP_000159368.1 | 6 | |
| | | WP_001288192.1 | 7 | |
| | | WP_000774603.1 | 8 | |
| | | WP_001001959.1 | 9 | |
| | | WP_000879943.1 | 10 | |
| *Streptococcus agalactiae* strain MRI Z1-022 | NZ_ANEM01000074.1 | WP_017648179.1 | 1 | |
| | | WP_017648177.1 | 2 | |
| | | WP_000802599.1 | 3 | |
| | | WP_025195242.1 | 4 | |

| | | | | |
|---|---|---|---|---|
| *Streptococcus agalactiae* strain DK-NI-014 | NZ_CQEL01000002.1 | WP_011058321.1 | 1 | ML10 |
| | | WP_000965642.1 | 2 | |
| | | WP_000906736.1 | 3 | |
| | | WP_000076715.1 | 4 | |
| | | WP_001183891.1 | 5 | |
| | | WP_000165747.1 | 6 | |
| | | WP_000763909.1 | 7 | |
| | | WP_000166038.1 | 8 | |
| | | WP_001107539.1 | 9 | |
| | | WP_001001742.1 | 10 | |
| *Streptococcus agalactiae* strain DK-PW-096 | NZ_MAWX01000026.1 | WP_000258802.1 | 1 | |
| | | WP_001921522.1 | 2 | |
| | | WP_000774601.1 | 3 | |
| | | WP_000218309.1 | 4 | |
| | | WP_000411527.1 | 5 | |
| | | WP_001270064.1 | 6 | |
| | | WP_000659174.1 | 7 | |
| | | WP_001042771.1 | 8 | |
| | | WP_000609259.1 | 9 | |
| | | WP_000071276.1 | 10 | |
| *Streptococcus agalactiae* strain GB-NI-015 | NZ_CTJD01000013.1 | WP_000660738.1 | 1 | |
| | | WP_017827941.1 | 2 | |
| | | WP_000965655.1 | 3 | |
| | | WP_000138374.1 | 4 | |
| | | WP_001872365.1 | 5 | |
| | | WP_000614971.1 | 6 | |
| | | WP_000258802.1 | 7 | |
| | | WP_000763911.1 | 8 | |
| | | WP_000118546.1 | 9 | |
| | | WP_000239531.1 | 10 | |
| *Streptococcus agalactiae* strain IT-NI-009 | NZ_CPZS01000003.1 | WP_079261174.1 | 1 | |
| | | WP_050201842.1 | 2 | |
| | | WP_000138374.1 | 3 | |
| | | WP_000474006.1 | 4 | |
| | | WP_000258802.1 | 5 | |
| | | WP_047200447.1 | 6 | |

| | | | |
|---|---|---|---|
| | | WP_000695684.1 | 7 |
| | | WP_047200442.1 | 8 |
| | | WP_001921522.1 | 9 |
| | | WP_047200443.1 | 10 |
| *Streptococcus agalactiae* strain RBH12 | NZ_CPVQ01000026.1 | WP_000650503.1 | 1 |
| | | WP_050198474.1 | 2 |
| | | WP_001058281.1 | 3 |
| | | WP_079454162.1 | 4 |
| | | WP_050199334.1 | 5 |
| | | WP_000612386.1 | 6 |
| | | WP_000963485.1 | 7 |
| | | WP_000206191.1 | 8 |
| | | WP_000461517.1 | 9 |
| | | WP_050199335.1 | 10 |
| *Streptococcus agalactiae* strain ST 618 | NDGB01000030.1 | OTG45475.1 | 1 |
| | | OTG45477.1 | 2 |
| | | OTG45479.1 | 3 |
| | | OTG45481.1 | 4 |
| | | OTG45480.1 | 5 |
| | | OTG45476.1 | 6 |
| | | OTG45474.1 | 7 |
| | | OTG45478.1 | 8 |
| | | OTG45473.1 | 9 |
| | | OTG45746.1 | 10 |
| *Streptococcus oralis* strain DD14 | NZ_KQ969340.1 | WP_061420077.1 | 1 |
| | | WP_061420097.1 | 2 |
| | | WP_061420111.1 | 3 |
| | | WP_061420115.1 | 4 |
| | | WP_061420080.1 | 5 |
| | | WP_061420123.1 | 6 |
| | | WP_061420133.1 | 7 |
| | | WP_061420073.1 | 8 |
| | | WP_061420062.1 | 9 |
| | | WP_061420131.1 | 10 |
| | NZ_KQ961462.1 | WP_061100257.1 | 1 |
| | | WP_061100237.1 | 2 |

| | | | |
|---|---|---|---|
| *Streptococcus pasteurianus* strain GED7275A | | WP_061100224.1 | 3 |
| | | WP_061100243.1 | 4 |
| | | WP_061100244.1 | 5 |
| | | WP_061100238.1 | 6 |
| | | WP_061100250.1 | 7 |
| | | WP_061100226.1 | 8 |
| | | WP_061100248.1 | 9 |
| | | WP_048791342.1 | 10 |
| *Streptococcus pyogenes* GA03805 | NZ_AWTL01000007.1 | WP_023079933.1 | 1 |
| | | WP_023079900.1 | 2 |
| | | WP_023079918.1 | 3 |
| | | WP_002985387.1 | 4 |
| | | WP_011528776.1 | 5 |
| | | WP_023079897.1 | 6 |
| | | WP_011017565.1 | 7 |
| | | WP_023079923.1 | 8 |
| | | WP_023079938.1 | 9 |
| | | WP_011888757.1 | 10 |
| *Streptococcus pyogenes* strain SST2091-1 | NZ_LRGN01000012.1 | WP_011889039.1 | 1 |
| | | WP_011285632.1 | 2 |
| | | WP_010922455.1 | 3 |
| | | WP_010922464.1 | 4 |
| | | WP_002994744.1 | 5 |
| | | WP_063629031.1 | 6 |
| | | WP_063629030.1 | 7 |
| | | WP_063629029.1 | 8 |
| | | WP_063629037.1 | 9 |
| | | WP_063629028.1 | 10 |
| *Streptococcus pyogenes* strain SST2097-1 | LRGT01000330.1 | OAC70939.1 | 1 |
| | | OAC70918.1 | 2 |
| | | OAC70928.1 | 3 |
| | | OAC70915.1 | 4 |
| | | OAC70921.1 | 5 |
| | | OAC70938.1 | 6 |
| | | OAC70937.1 | 7 |
| | | OAC70919.1 | 8 |

| | | | | |
|---|---|---|---|---|
| | | OAC70916.1 | 9 | |
| | | OAC70925.1 | 10 | |
| *Listeria monocytogenes* strain R2-502 | NC_021838.1 | WP_003731277.1 | 1 | ML2 |
| | | WP_003731276.1 | 2 | ML3 |
| | | WP_023553812.1 | 3 | AcrIIA1 |
| | | WP_003733721.1 | 4 | |
| | | WP_003731655.1 | 5 | |
| | | WP_014601388.1 | 6 | |
| | | WP_023553826.1 | 7 | |
| | | WP_003722567.1 | 8 | |
| | | WP_003730996.1 | 9 | |
| | | WP_003731704.1 | 10 | |
| | | WP_023553814.1 | 16 | AcrIIA2 |
| | | WP_003722518.1 | 38 | AcrIIA1 |
| | | WP_012581438.1 | 56 | AcrIIA1 |
